# scTGCL: A Transformer-Based Graph Contrastive Learning Approach for Efficiently Clustering Single-Cell RNA-seq Data

**DOI:** 10.64898/2026.03.28.714542

**Authors:** Md. Shoaib Abdullah Khan, Md. Mahir Faisal, Md. Humayun Kabir

## Abstract

**Motivation:** Single-cell RNA sequencing (scRNA-seq) enables characterization of cellular heterogeneity at unprecedented resolution. However, accurate clustering of scRNA-seq data remains challenging due to high dimensionality, dropout-induced sparsity, and technical noise. Existing contrastive learning methods usually create augmented views by perturbing gene expression values, so the uncertainty in cell-cell relationships caused by dropout events is not properly modeled.

**Results:** We propose scTGCL, a novel Transformer-based graph contrastive learning framework that learns robust and discriminative cell representations directly from gene expression data. scTGCL employs multi-head self-attention to adaptively construct weighted cell-cell graphs and a Transformer encoder to generate cell embeddings from them, capturing diverse biological relationships. For contrastive augmentation, we apply feature-level gene masking together with attention masking that randomly drops edges of the learned attention matrices, enabling the model to withstand dropout events and technical noise. A symmetric contrastive loss maximizes agreement between original and augmented representations, while joint optimization with reconstruction and imputation losses preserves biological interpretability. Extensive experiments on twenty real and simulated scRNA-seq datasets demonstrate that scTGCL achieves the highest average performance among nine state-of-the-art baseline methods in clustering accuracy, normalized mutual information, and adjusted Rand index, while also achieving superior imputation performance and substantially lower runtime on large-scale datasets. Furthermore, biological analysis confirms that the learned clusters align with known cell types and yield biologically interpretable pathway enrichment, validating the utility of scTGCL for downstream cell annotation and marker gene discovery.

**Availability and Implementation:** The source code and test data are freely available at GitHub (https://github.com/ShoaibAbdullahKhan/scTGCL).

## 1. Introduction

Cells serve as the fundamental structural and functional units of all living organisms. Understanding cellular behavior through transcriptomic profiling is essential for advancing biological research and medical applications [1]. Conventional techniques such as microarrays and bulk RNA sequencing measure gene expression as an average signal across a large population of cells, thereby masking cell-to-cell variability. The advent of single-cell RNA sequencing (scRNA-seq) technologies has addressed this limitation by enabling gene expression analysis at the resolution of individual cells, allowing researchers to uncover cellular heterogeneity and transcriptional diversity that are otherwise unobservable in bulk measurements [2].

A core objective in scRNA-seq data analysis is the grouping of cells with similar transcriptional profiles into meaningful clusters. Accurate clustering facilitates the identification of distinct cell types and functional states and supports a wide range of downstream analyses. However, clustering scRNA-seq data remains a challenging task due to its intrinsic characteristics, including high dimensionality, extensive sparsity caused by dropout events, and technical noise [3, 4]. These issues complicate the accurate estimation of cell–cell similarity and hinder the performance of conventional computational methods. Furthermore, the rapidly increasing scale of modern scRNA-seq datasets introduces additional computational burdens that demand more robust and scalable analytical frameworks.

To address these challenges, deep embedded clustering methods have been extensively explored to jointly learn low-dimensional representations and cluster assignments. Models such as scDCCA, scDeepCluster, DESC, scziDesk, scGMAI, scBGEDA, and DeepScena leverage autoencoder-based architectures combined with iterative or self-training clustering objectives to produce cluster-friendly latent spaces [5–11]. These approaches demonstrate strong scalability and improved clustering accuracy; however, many of them rely primarily on feature reconstruction and often overlook higher-order relational structures among cells.

To explicitly capture structural dependencies between cells, graph-based clustering frameworks have gained increasing attention. Methods such as GraphSCC, scDSC, scGAC, scDFC, scEGG, scLEGA, SCEA, scDFN, and scG-cluster model cells as nodes in a graph and exploit graph neural networks or graph attention mechanisms to encode cell–cell relationships [12–20]. By integrating structural and attribute information, these models significantly improve clustering robustness, though their performance remains sensitive to graph construction strategies. Moreover, most of these methods build the cell–cell graph once from a predefined similarity measure and keep it fixed during training, so noisy or missing connections caused by dropout events are passed directly into the learned cell embeddings.

More recently, contrastive learning has become a popular self-supervised strategy for scRNA-seq clustering. These methods create two views of each cell and train the model so that the two views of the same cell stay close in the embedding space, while views of different cells are pushed apart. Methods such as contrastive-sc, scNAME, and scRECL apply this idea to gene expression profiles [21–23]. Graph contrastive methods go a step further and also use a cell–cell graph. scGCL connects each cell to its *k* nearest neighbours by cosine similarity, scAGCL builds a *k*-nearest-neighbour graph from Pearson correlation and perturbs it to create its views, scMMN combines *k*-nearest-neighbour and diffusion-map graphs, and scGPCL builds a cell–gene graph from the expression matrix [24–27]. In all of these methods the graph is built once before training and then kept fixed. The weakness of a fixed graph is that it is computed from noisy and sparse expression values. When dropout events hide the expression of important genes, two cells of the same type can look dissimilar and two cells of different types can look similar, so wrong connections enter the graph and remain there for the whole of training. A graph that is learned and updated during training can correct these connections as the cell representations improve, and is therefore more reliable for clustering.

To avoid fixed graphs, a second group of methods uses multi-head attention to learn cell–cell relationships during training. AttentionAE-sc uses multi-head attention between cells to combine the embeddings of a denoising autoencoder and a graph autoencoder, but it does not use contrastive learning [28]. scSimGCL learns a cell–cell graph with a multihead attention module and combines it with graph contrastive learning [29]. However, it turns the attention scores into a binary graph by keeping only the scores above a threshold, so the strength of each relationship is lost, and the edges it drops for augmentation come from this binary graph rather than from the attention weights themselves. scSAMAC adds multi-head attention to a variational autoencoder and contrasts each cell with a view in which important genes are masked [30]. scHashFormer uses a Transformer to encode each cell together with neighbouring cells selected by hashing, and trains it with a contrastive loss that treats cells in the same hash bucket as positive pairs [31]. Although these methods learn cell–cell relationships dynamically, their augmentation is either absent or applied only to gene expression values or to a binary graph, and none of them masks the learned attention weights directly. This is an important gap for dropout handling. An attention weight records how much one cell relies on another, and when dropout distorts gene expression, some of these weights become unreliable. If we randomly remove part of the attention that a cell pays to other cells, the model can no longer depend on any single cell–cell connection and must learn representations that stay stable when some relationships are missing. In this way, attention masking directly models the uncertainty in cell–cell relationships caused by dropout, and it forms the core idea of our proposed model.

In addition, several hybrid and advanced frameworks have been proposed to further improve clustering accuracy and robustness. Models such as scAMAC, scMAE, scSemiAAE, scMSCF, scHFC, and scHetG integrate multi-scale attention mechanisms, masked autoencoders, semi-supervised learning, ensemble clustering, or heterogeneous graph modeling to capture complementary biological signals [32–37]. While these methods demonstrate promising performance, the increasing architectural complexity often leads to higher computational costs and reduced interpretability.

In this study, we propose scTGCL (single-cell Transformer-based Graph Contrastive Learning), a clustering framework for scRNA-seq data. scTGCL first projects the high-dimensional and sparse gene expression data into a lower-dimensional embedding, and then uses a Transformer-based encoder [38] with multi-head self-attention to learn how strongly each cell is related to the other cells. Each attention head therefore acts as a weighted cell–cell graph, and the encoder uses these relationships directly to generate the cell embeddings. The main novelty of scTGCL is an attention masking strategy for creating the augmented view used in contrastive learning. In this view, we randomly remove a portion of the attention weights between pairs of cells, so that the model loses access to some cell–cell relationships, and we also randomly mask a portion of the gene expression values. A contrastive loss then pulls together the representations of the original and augmented views of the same cell. Dropout events in scRNA-seq data hide true expression values and make some cell–cell relationships unreliable, so training the model to produce consistent representations when both genes and cell–cell connections are partly missing makes it more robust to dropout. To our knowledge, scTGCL is the first scRNA-seq clustering method to use attention masking in a Transformer encoder to create augmented views for contrastive learning. The contrastive loss is optimized together with reconstruction and imputation losses, so that the learned representation is both useful for clustering and able to recover missing expression values. Experiments on twenty real scRNA-seq datasets and simulated data show that scTGCL achieves strong clustering and imputation performance compared with existing methods.

## 2. Materials and methods

### 2.1. The framework of scTGCL

The overall structure of the proposed scTGCL, a contrastive Transformer-based autoencoder for scRNA-seq data analysis, is illustrated in Figure 1. The model is designed to learn robust and informative cell representations by jointly optimizing three key objectives: reconstruction, imputation, and contrastive learning. First, preprocessed gene expression data is passed through an encoder to create an initial embedding. This embedding is then fed into a multi-head attention module to learn a weighted cell-cell graph, capturing complex relationships between cells. Second, to create a robust augmented view for contrastive learning, we apply two distinct augmentation strategies: random gene masking on the input data and random edge dropping on the learned attention matrices. Third, both the original and augmented data are processed through a shared network to generate latent representations. Finally, a joint learning strategy is employed, combining a reconstruction loss, an imputation loss, and a contrastive loss.

**Fig 1.**
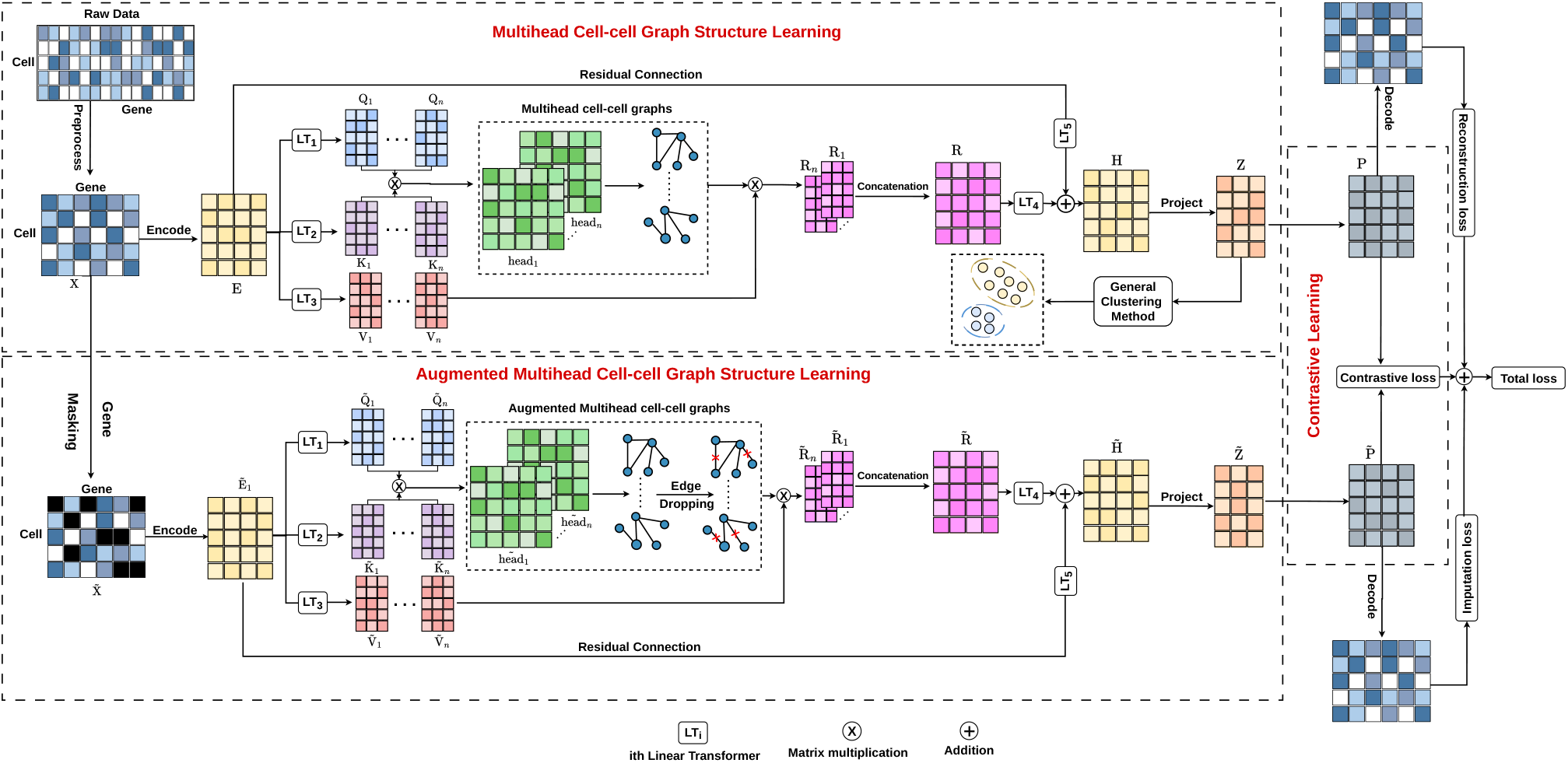
Architecture of the proposed scTGCL framework. The model learns cell–cell relations using multi-head attention, creates an augmented view by gene masking and edge dropping, and jointly optimizes reconstruction, imputation and contrastive losses.

### 2.2. Data Collection and Preprocessing

We evaluated the proposed model on twenty real scRNA-seq datasets collected from diverse studies spanning multiple tissues, species, and sequencing technologies. These datasets cover both human and mouse tissues across a broad range of organs including brain, pancreas, liver, retina, kidney, lung, testis, mammary gland, muscle, and blood, and encompass diverse biological conditions and cell populations. The number of cells ranges from 90 to 54,865, the number of genes ranges from 4,999 to 33,694, and the number of cell types ranges from 4 to 55, reflecting substantial variation in dataset scale and complexity. The datasets were generated using several sequencing platforms including 10x Chromium, InDrop, SMARTer, STRT-seq UMI, Drop-seq, CEL-seq2, Smart-seq2, and the Tang protocol. All datasets provide ground-truth cell-type labels and have been widely used in previous clustering studies, making them suitable for rigorous performance comparison. Detailed statistics of all datasets are summarized in Table 1.

**Table 1.**
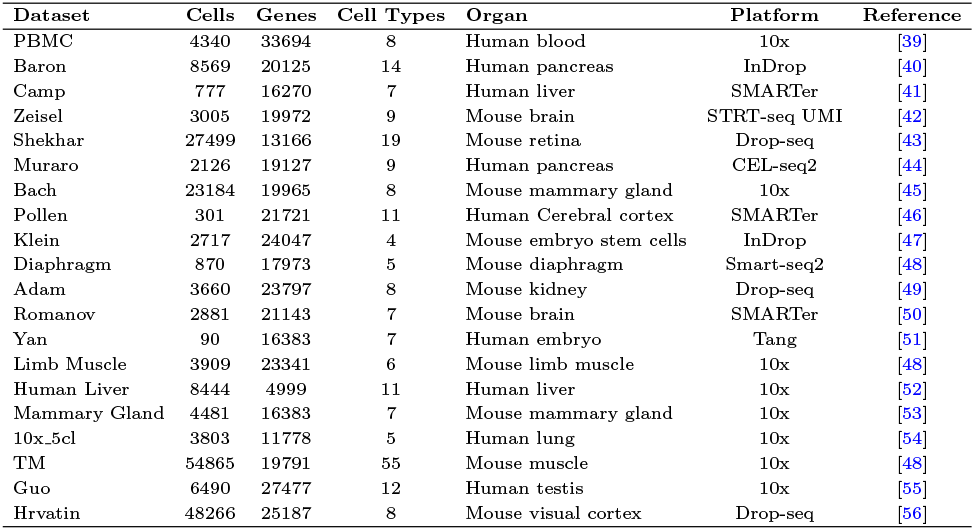
Summary of scRNA-seq datasets used in this study.

Each dataset is represented as a gene expression matrix where each row contains the expression values of all genes for a particular cell. Before training, the data is preprocessed using the Python Scanpy package [57]. Genes not expressed in any cell are first removed, after which 2500 highly variable genes (HVGs) are selected to retain the most informative features. Expression values are then normalized by scaling each cell to a total count of 10^4^, and a log transformation is applied to reduce the effect of extreme values and stabilize the data distribution.

### 2.3. Multi-head Cell-Cell Graph Structure Learning

The first core component of scTGCL is the multi-head cell-cell graph structure learning module. The primary goal of this module is to transform the raw, high-dimensional gene expression matrix into a lower-dimensional latent space while simultaneously learning a weighted graph that captures the intricate similarities and relationships between cells.

Let *X* ∈ ℝ^*N* ×*g*^ be the pre-processed gene expression matrix, where *N* is the number of cells in a batch and *g* is the number of genes. The process begins by projecting this high-dimensional input into a fixed-dimensional embedding space.

This is achieved through an input projection layer that maps each cell’s gene expression profile to a dense embedding vector.

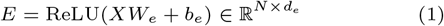

Where 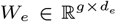 and 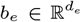 are learnable parameters of the projection layer, *d*_*e*_ is the embedding dimension, and the bias is broadcast across the *N* rows.

This embedding matrix *E* serves as the input to a multi-head attention module, which is responsible for learning the cell-cell graph. The attention mechanism allows the model to dynamically weigh the influence of all other cells on a given cell. For each attention head *i* (where *i* ∈ {1, …, *n*}), we first generate query (*Q*_*i*_), key (*K*_*i*_), and value (*V*_*i*_) matrices by applying linear transformations to the embedding *E*.

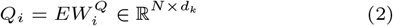

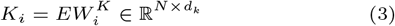

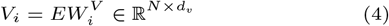

Where 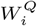, 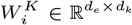 and 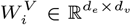 are learnable weight matrices for the *i*-th head.

The cell-cell graph for a single head is derived by computing the dot product similarity between the query and key matrices. A softmax function is then applied row-wise to normalize these scores into a probability distribution, yielding the weighted adjacency matrix, or attention head, *head*_*i*_.

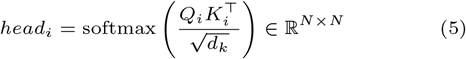

Each head *head*_*i*_ can be interpreted as a weighted, fully-connected cell-cell graph. The weight *head*_*i*_[*j, k*] signifies the importance or relevance of cell *k* to cell *j* from the perspective of the *i*-th attention head. Different heads can capture different types of biological relationships, such as similarities based on a specific subset of genes.

To generate meaningful feature vectors that incorporate these relational graphs, we multiply each attention head *head*_*i*_ by its corresponding value matrix *V*_*i*_. This operation aggregates information from neighboring cells according to the learned graph structure, producing an output *R*_*i*_ for each head.

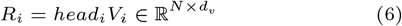

These head-specific outputs, *R*_1_ … *R*_*n*_, capture distinct, relation-aware features of the cells. They are then concatenated along the feature dimension to form a single comprehensive matrix 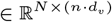.

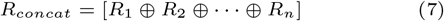

This concatenated representation is passed through a final linear transformation (*W*_*O*_), while the original input embedding *E* is simultaneously projected through a separate learnable linear transformation (*W*_*I*_) along the residual path. The two projected signals are then summed and passed through layer normalization:

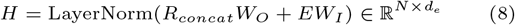

Where 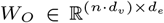 and 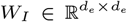. By projecting both the attention output and the residual through dedicated weight matrices, the model can learn the optimal contribution of each pathway rather than adding the raw embedding as a fixed skip connection.

The resulting matrix (*H*) is a robust, graph-aware embedding of the cells. The latent representation 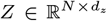 is then obtained by projecting *H* into a lower-dimensional space using an encoder output layer. This lower-dimensional representation is the core embedding used for downstream tasks like clustering.

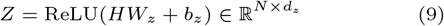

where 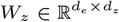 and 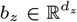.

Finally, to ensure that this latent space retains the essential information from the original data, a decoder reconstructs the gene expression matrix 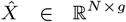 from the latent representation *Z*.

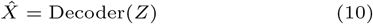

The fidelity of this reconstruction is measured by the reconstruction loss, which is the mean squared error (MSE) between the original data and the reconstructed output.

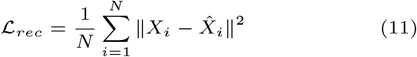

This loss function ensures that the latent representation *Z* preserves the global structure and key patterns of the original gene expression data.

### 2.4. Augmented Multi-head Cell-Cell Graph Structure Learning

To learn representations that are robust to noise and technical artifacts common in scRNA-seq data, we introduce an augmented view of the data. This view is generated through a two step augmentation strategy applied within the same multi-head graph learning framework. The model is then trained to pull the representations of the original and augmented views of the same cell closer together, while pushing apart representations of different cells.

First, an augmented gene expression matrix,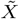, is created by randomly masking a subset of gene expression values to zero for each cells. This is a form of feature augmentation that simulates the effect of dropout events. A binary keep-mask *M* ^(*x*)^ ∈ {0, 1}^*N* ×*g*^ is generated, where 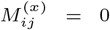 indicates that the expression of gene *j* in cell *i* is masked to zero. The augmented matrix is then created by element-wise multiplication.

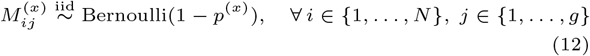

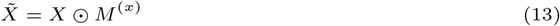

The mask has the same shape as the full expression matrix, and every cell–gene entry (*i, j*) receives its own independent Bernoulli draw. Consequently, each cell *i* is assigned a different randomly selected subset of masked genes, and no gene is masked across all cells simultaneously unless this occurs by chance. The genes to be masked are chosen uniformly at random, so that on average a fraction *p*^(*x*)^ of the measured entries of each cell is hidden. Moreover, *M* ^(*x*)^ is resampled independently at every training step, so a given cell is exposed to a different set of masked genes at each epoch.

This augmented matrix 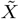 is then passed through the same encoder and multi-head attention module to produce a new set of attention heads, 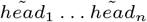. At this stage, we apply a second augmentation technique: graph augmentation via edge dropping. This involves randomly setting a fraction of the weights in each learned attention head 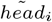 to zero. This simulates uncertainty in the cell-cell relationships and forces the model to learn from a sparser, more robust graph structure. For every head *i*, a separate mask *M* ^(*a,i*)^ ∈ {0, 1}^*N* ×*N*^ is generated, where 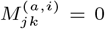 removes the edge between cell *j* and cell *k* in that head. Crucially, the diagonal (self-attention) is kept intact to preserve the identity of each cell, and each row is re-normalized after masking so that the surviving attention weights of every cell again sum to one.

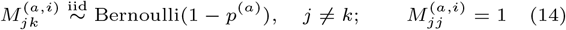

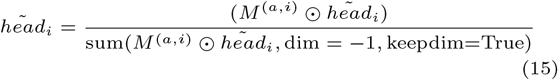

As with gene masking, the edge mask is not shared. An independent mask *M* ^(*a,i*)^ is drawn for each of the *n* attention heads, and within each head every off-diagonal entry (*j, k*) is sampled independently. Therefore each cell *j* loses its own random subset of neighbours, this subset differs from head to head, and the same cell pair may be dropped in one head while being retained in another. The edges to be dropped are selected uniformly at random, so that on average a fraction *p*^(*a*)^ of the off-diagonal edges of each cell is removed in each head. The masks are likewise resampled at every training step. Because the heads are perturbed independently, the model cannot rely on any single cell–cell connection and must recover the cell relationships from the consensus of the remaining edges across heads.

The rest of the pipeline, including the residual connection, encoder output layer, and decoder, is applied to this augmented view, resulting in an augmented latent representation 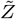 and a reconstructed output 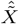. This reconstruction from the augmented data gives rise to the imputation loss. The imputation loss measures the model’s ability to recover the original, non-masked data from its corrupted, augmented version. This task forces the model to learn the underlying data manifold and fill in missing values, acting as a powerful regularizer.

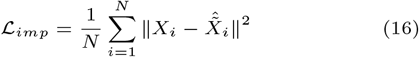

### 2.5. Contrastive Learning

The final component of the framework is the contrastive learning module, which operates on the latent representations *Z* and 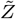. Before computing the contrastive loss, these representations are passed through a small projection head, a multilayer perceptron that maps them to a space where the contrastive loss is applied yielding *P* = Proj(*Z*) and 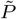 = Proj 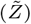.

The contrastive loss is designed to maximize the agreement between the two augmented views of the same cell (positive pairs) while minimizing the agreement between views of different cells (negative pairs). For a given cell *i*, its representation from the original view *P*_*i*_ and from the augmented view 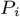 form the positive pair. All other samples in the batch are treated as negative examples.

To compute the loss, the embeddings are first normalized to have unit length. Three similarity matrices are then computed using dot products, scaled by a temperature parameter *τ*. These matrices represent the pairwise similarities between all cells in the original view *S*^oo^, the augmented view *S*^aa^, and across the two views *S*^oa^.

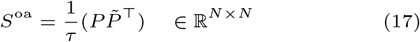

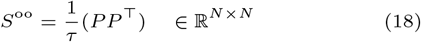

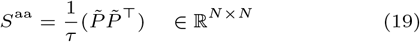

The positive similarity for cell *i* is defined as the similarity between its original and augmented representation, which corresponds to the diagonal element of the cross-view similarity matrix.

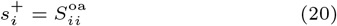

For the loss computed from the original view to the augmented view, the exponential of the similarities are taken to obtain positive weights. The denominator for a given cell *i* consists of two parts: (i) the sum of its similarities to all cells in the augmented view 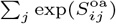, and (ii) the sum of its similarities to all other cells in its own (original) view 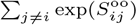. The diagonal of the original-to-original similarity matrix is explicitly masked out to prevent a cell from treating itself as a negative sample.

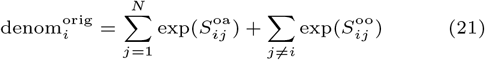

The loss for this direction is then the negative log probability of the positive pair.

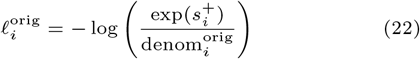

A symmetric loss is computed from the augmented view to the original view. Here, the denominator for cell *i* is the sum of its similarities to all cells in the original view 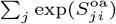 and the sum of its similarities to all other cells in its own augmented view 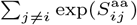.

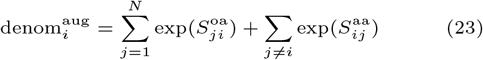

The corresponding loss is:

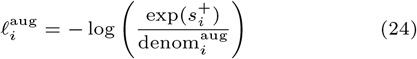

The final contrastive loss is the average of these two symmetric losses over all cells in the batch.

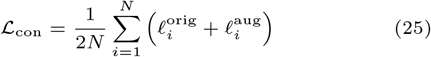

This contrastive objective encourages the encoder to be invariant to the applied augmentations, making the learned representations more robust and discriminative.

### 2.6. Joint Optimization

The overall training objective of scTGCL is to jointly minimize the three loss functions described above. The total loss is a weighted sum of the reconstruction, imputation, and contrastive losses.

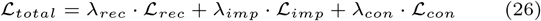

The hyperparameters *λ*_*rec*_, *λ*_*imp*_, and *λ*_*con*_ balance the contribution of each component during training. By optimizing this joint loss function, scTGCL learns latent cell representations that are simultaneously faithful to the original data, robust to noise and capable of imputing missing values, and highly discriminative for separating different cell types. The final representation *Z* can then be used for downstream analyses such as clustering or visualization, as evaluated in the results section.

### 2.7. Implementation and parameters setting

The proposed scTGCL is implemented using Python 3.8+ with the PyTorch (version 1.9+) deep learning framework. For each dataset, the preprocessed expression matrix is split into 80% for model training and 20% for testing, with stratification by cell type to preserve the class proportions. The input projection layer maps each cell’s expression profile through a hidden layer of 2048 units into an embedding of dimension *d*_*e*_ = 512, on which a multi-head self-attention module with 8 attention heads learns the cell–cell graph. The resulting graph-aware representation is compressed into a *d*_*z*_ = 64 dimensional latent space, projected to a 256-dimensional contrastive space, and reconstructed by a symmetric decoder, with a dropout rate of 0.3 applied at the input projection to mitigate overfitting.

For contrastive augmentation, the gene masking probability and the attention masking probability are set to *α* = 0.25 and *β* = 0.35, respectively. The overall objective is a weighted sum of the reconstruction, imputation, and contrastive losses with weights *λ*_*rec*_ = 6, *λ*_*imp*_ = 1, and *λ*_*con*_ = 0.8, and the contrastive temperature is fixed at *τ* = 0.5. The network is optimized using the AdamW optimizer with a learning rate of 1 × 10^−3^ and a weight decay of 1 × 10^−3^, and the learning rate is annealed following a cosine schedule over the course of training. Gradients are clipped to a maximum *L*_2_ norm of 1.0 to stabilize optimization, and each model is trained for 200 epochs. The batch size is adapted to the dataset scale: 128 for datasets with fewer than 4000 cells, 512 for those with 4000 to 8000 cells, and 1024 for datasets exceeding 8000 cells.

The learned latent embeddings are clustered using the K-Means algorithm from Scikit-learn with 20 centroid initializations, where the number of clusters is set to the known number of cell types in each dataset. To ensure reproducibility, all random number generators of Python, NumPy, and PyTorch are seeded and deterministic cuDNN operations are enforced; all reported metrics are averaged over ten independent runs with different random seeds. These hyperparameter settings are obtained through grid search together with the sensitivity analysis reported in Supplementary Figure S2. All experiments are conducted on a workstation equipped with an AMD Ryzen 9 7900X CPU, 32 GB of RAM, and an NVIDIA RTX 5070 Ti GPU.

### 2.8. Clustering and Evaluation Metrics

To evaluate the clustering performance of the proposed scTGCL framework, we employ three widely used clustering metrics: clustering accuracy (CA), normalized mutual information (NMI), and adjusted Rand index (ARI). These metrics provide a comprehensive assessment of the similarity between predicted clustering assignments and the ground-truth cell-type annotations. After training, the encoder generates a latent embedding matrix *Z*, which captures biologically meaningful and noise-resilient representations of individual cells. These embeddings are subsequently clustered using general-purpose algorithms such as K-Means or Leiden to produce predicted cluster labels *Ŷ*. Let *Y* denote the ground-truth cell-type labels and *N* the total number of cells in the dataset.

#### 2.8.1. CA

The clustering accuracy measures the proportion of correctly matched cells between the predicted and ground-truth clusters. It is defined as:

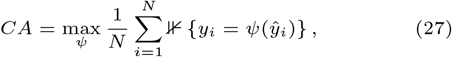

where *ψ* represents a one-to-one mapping function that aligns predicted clusters with true cell types, and ⊮{·} is an indicator function. A higher CA value indicates better alignment between predicted and actual biological labels.

#### 2.8.2. NMI

The normalized mutual information quantifies the shared information between the predicted cluster assignments 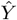 and the ground-truth labels *Y*. It is expressed as:

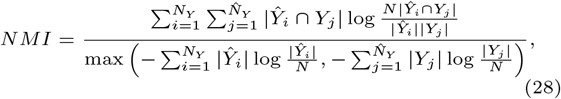

where 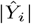 and |*Y*_*j*_ | denote the number of cells in the *i*-th predicted cluster and the *j*-th ground-truth cluster, respectively. NMI values range between 0 and 1, with higher values indicating stronger consistency between predicted and true clusters.

#### 2.8.3. ARI

The adjusted Rand index measures the similarity between two cluster assignments by considering all pairs of cells and determining whether they are consistently assigned to the same or different clusters in both *Y* and 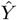. The ARI is calculated as follows:

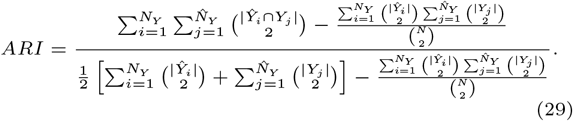

The ARI value ranges from −1 to 1, where 1 denotes perfect agreement, 0 indicates random labeling, and negative values imply disagreement between clusterings.

In addition to clustering metrics, we assess the imputation capability of scTGCL using two complementary metrics: L1 distance and Pearson correlation coefficient (PCC). Let *X* ∈ ℝ^*N* ×*G*^ denote the original gene expression matrix with *N* cells and *G* genes, and let 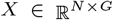 denote the corresponding imputed matrix.

#### 2.8.4. L1 Distance

The L1 distance measures the average absolute difference between the imputed and original expression values across all cells and genes:

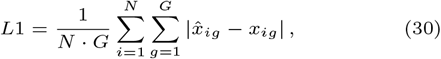

where 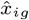 and 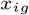 denote the imputed and original expression value of gene *g* in cell *i*, respectively. A lower L1 distance indicates more accurate recovery of the original expression profile.

#### 2.8.4. PCC

The Pearson correlation coefficient measures the linear correspondence between the imputed and original expression profiles. For each cell *i*, the per-cell PCC is computed as:

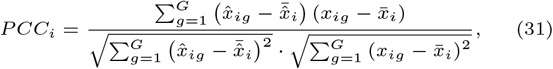

where 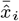 and 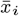 are the mean imputed and mean original expression values across all genes in cell *i*, respectively. The overall PCC is reported as the average over all cells:

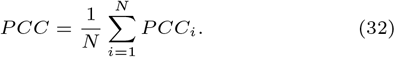

PCC values range from −1 to 1, with higher values indicating stronger agreement between the imputed and original expression patterns.

## 3. Results and Discussion

### 3.1. Performance comparison with baseline models

In this study, we evaluate the clustering performance of the proposed scTGCL framework on twenty real scRNA-seq datasets and compare it with nine representative baseline methods: scSimGCL [29], scMAE [33], scAGCL [25], CIDR [58], scDCCA [5], k-means [59], scMMN [26], scAMAC [32], scDFN [19]. Each model is run independently ten times on each dataset, and the averaged results of three clustering metrics: CA, NMI, and ARI are reported to assess overall performance.

Figure 2 compares the clustering performance of all methods on the twenty datasets, with CA shown in Figure 2(A), NMI in Figure 2(B) and the ARI results are given in Supplementary Figure S1. Each bar is the average of ten runs with different random seeds, and the error bar on top of it shows the standard deviation across these runs, so the stability of each method can be compared alongside its accuracy. scTGCL achieves the best score on 15 of the 20 datasets for both CA and NMI and on 14 datasets for ARI. Averaged over all twenty datasets, scTGCL also obtains the highest CA (84.98%), NMI (87.13%), and ARI (80.47%), ahead of the next best method, scSimGCL (80.50%, 83.12%, and 75.13%). On the datasets where scTGCL is not the best, it is usually behind by only a small margin.

**Fig 2.**
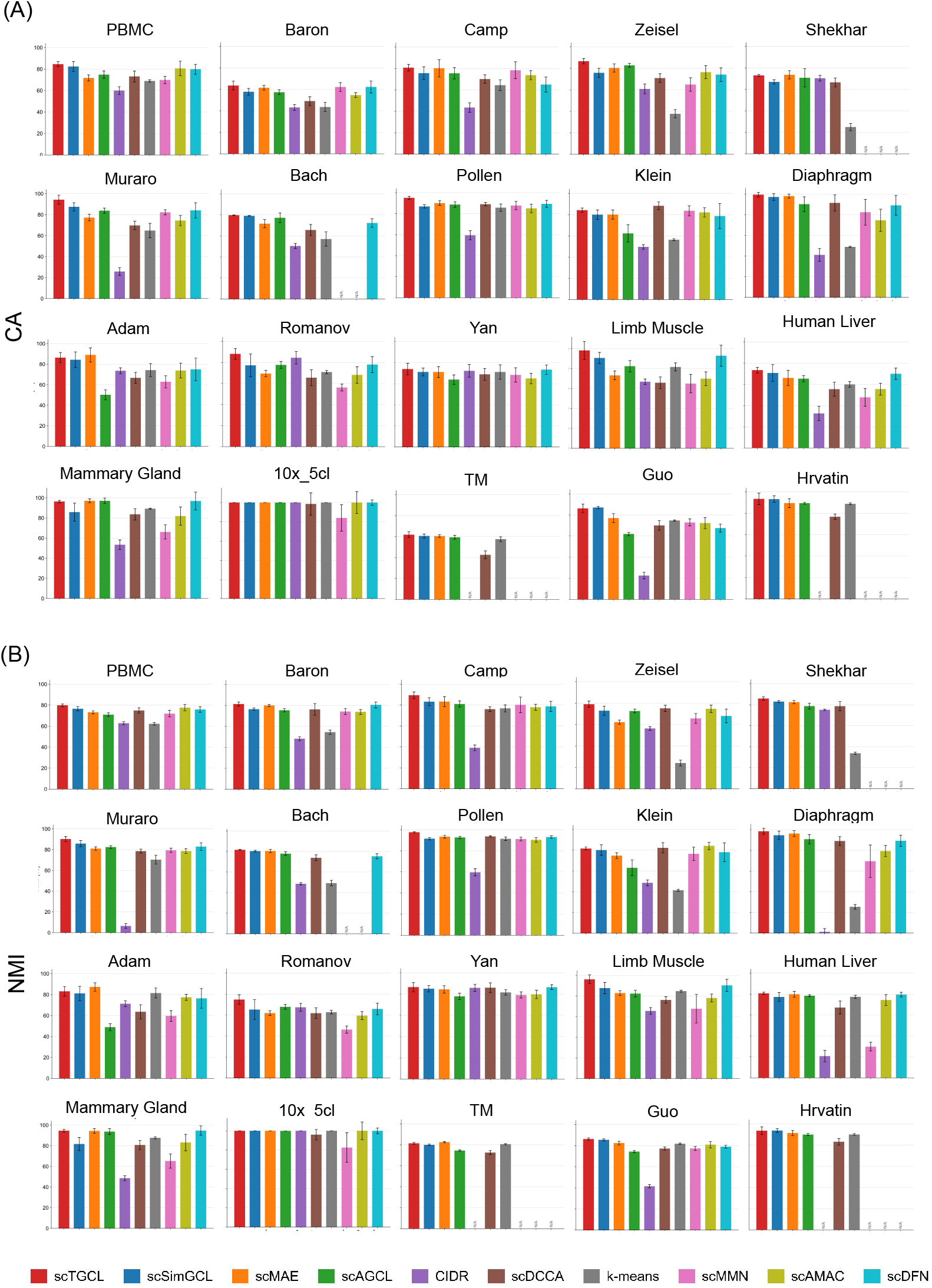
Clustering performance of scTGCL and nine baseline methods on twenty real scRNA-seq datasets. (A) Clustering accuracy (CA, %). (B) Normalized mutual information (NMI, %). Each bar shows the mean over ten independent runs with different random seeds, and the error bar on top of it shows the standard deviation across these runs. N/A marks a method that could not be run on that dataset. The corresponding ARI results are shown in Supplementary Figure S1.

Compared with traditional methods such as k-means and CIDR, scTGCL shows large improvements on all three metrics, highlighting the importance of learning expressive latent representations rather than relying on handcrafted similarity measures. Deep graph and contrastive baselines such as scSimGCL and scMAE are more competitive, but scTGCL still improves on them on most datasets, which we attribute to its adaptive attention-based cell–cell graphs and to the attention masking augmentation that makes the learned representations robust to dropout. Some baselines could not be run on the largest datasets: CIDR failed on TM and Hrvatin, scDFN on Shekhar, TM, and Hrvatin, and scMMN and scAMAC on Shekhar, Bach, TM, and Hrvatin.

To further illustrate the intuitive clustering quality, we visualize the learned embeddings using t-SNE on two representative datasets with different scales and cell-type complexities, namely Baron and Muraro. Figure 3 shows the two-dimensional t-SNE projections of embeddings generated by all compared methods, where Figure 3(A) corresponds to the Baron dataset and Figure 3(B) corresponds to the Muraro dataset.

**Fig 3.**
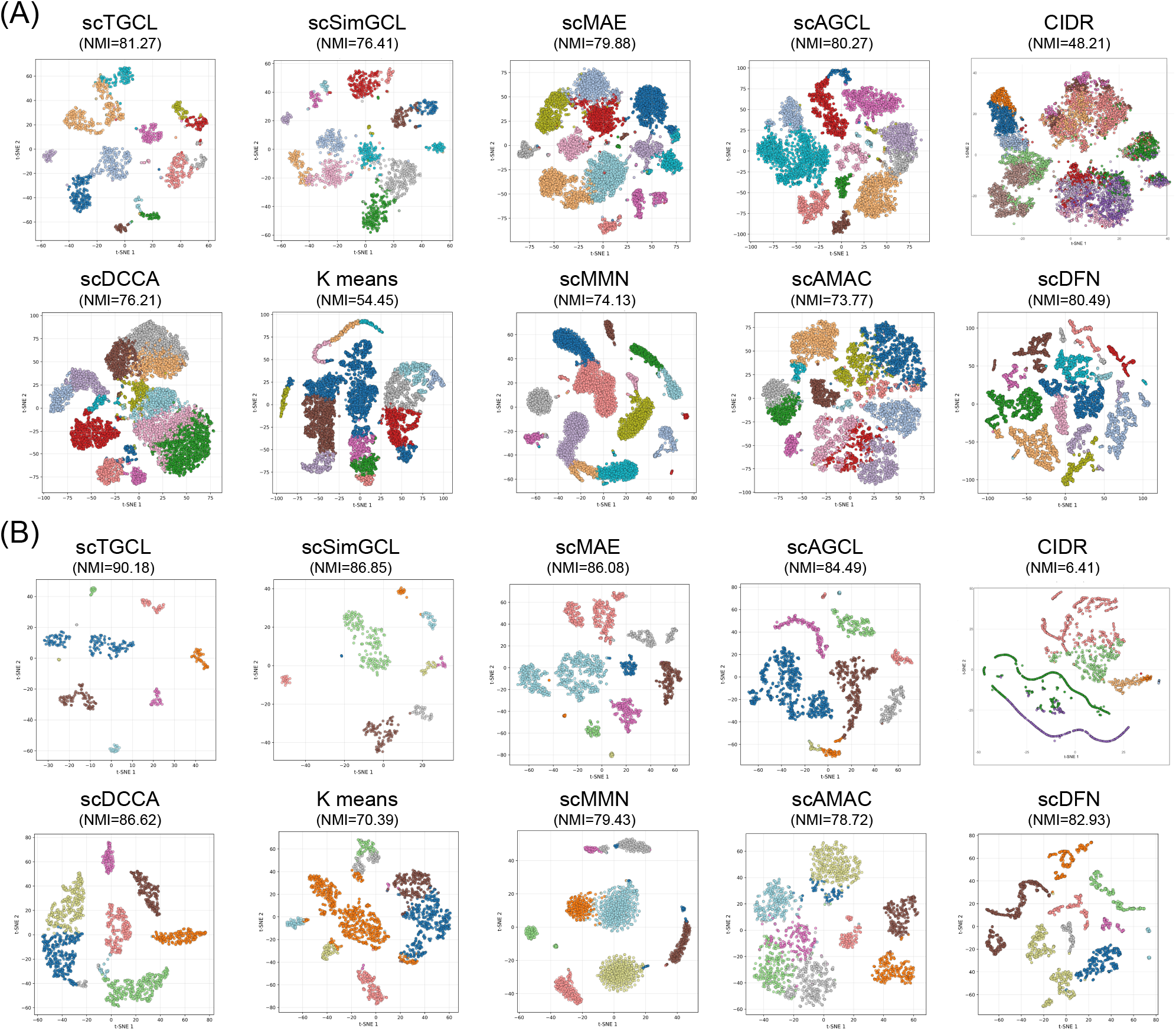
t-SNE visualization of cell embeddings learned by different methods on (A) Baron and (B) Muraro datasets. Colors denote ground-truth cell types. scTGCL produces more compact intra-cluster structures and clearer inter-cluster boundaries compared with baseline methods.

As shown in Figure 3, scTGCL yields well-separated and compact clusters on both datasets, with minimal overlap between different cell types. On the Baron dataset, scTGCL produces clearer cluster boundaries compared with other methods, many of which exhibit substantial mixing between cell subpopulations. On the Muraro dataset, scTGCL further demonstrates its ability to preserve fine-grained cellular structure, resulting in embeddings that closely align with the ground-truth annotations. In contrast, several baseline models fail to adequately separate rare or closely related cell types, leading to ambiguous cluster assignments.

To further evaluate the capability of scTGCL to recover missing gene expression values caused by dropout events, we conduct an imputation experiment on all twenty real scRNA-seq datasets. Artificial dropout events are introduced into the gene expression matrix at three rates: 10%, 20%, and 30%, and the ability of each model to reconstruct the original expression profiles is assessed using two complementary metrics: L1 distance, where lower values indicate more accurate recovery, and Pearson correlation coefficient (PCC), where higher values reflect stronger agreement with the ground-truth expression patterns. Both metrics are computed only on the entries that were artificially masked: for these positions, we compare the values reconstructed by each model with the original values before masking. In this way, each method is judged solely on how well it recovers expression values it could not see, and the unmasked entries, which a model can reproduce simply by copying its input, do not affect the results (Figure 4). We compare scTGCL against four representative scRNA-seq imputation methods: GE-Impute [60], scGCL [24], scGNN [61], and AutoClass [62]. GE-Impute leverages graph embedding techniques to model cell-cell similarity and recover dropout entries in the expression matrix. scGCL integrates graph contrastive learning with a Zero-inflated Negative Binomial distribution to account for the stochastic nature of scRNA-seq data and impute missing values. scGNN employs a graph neural network framework that jointly infers cell-cell relationships and imputes dropout events through iterative feature aggregation. AutoClass combines an autoencoder with a classifier network, performing imputation as part of a broader deep learning pipeline for single-cell analysis. All methods are executed ten times on each dataset and the averaged results are reported.

**Fig 4.**
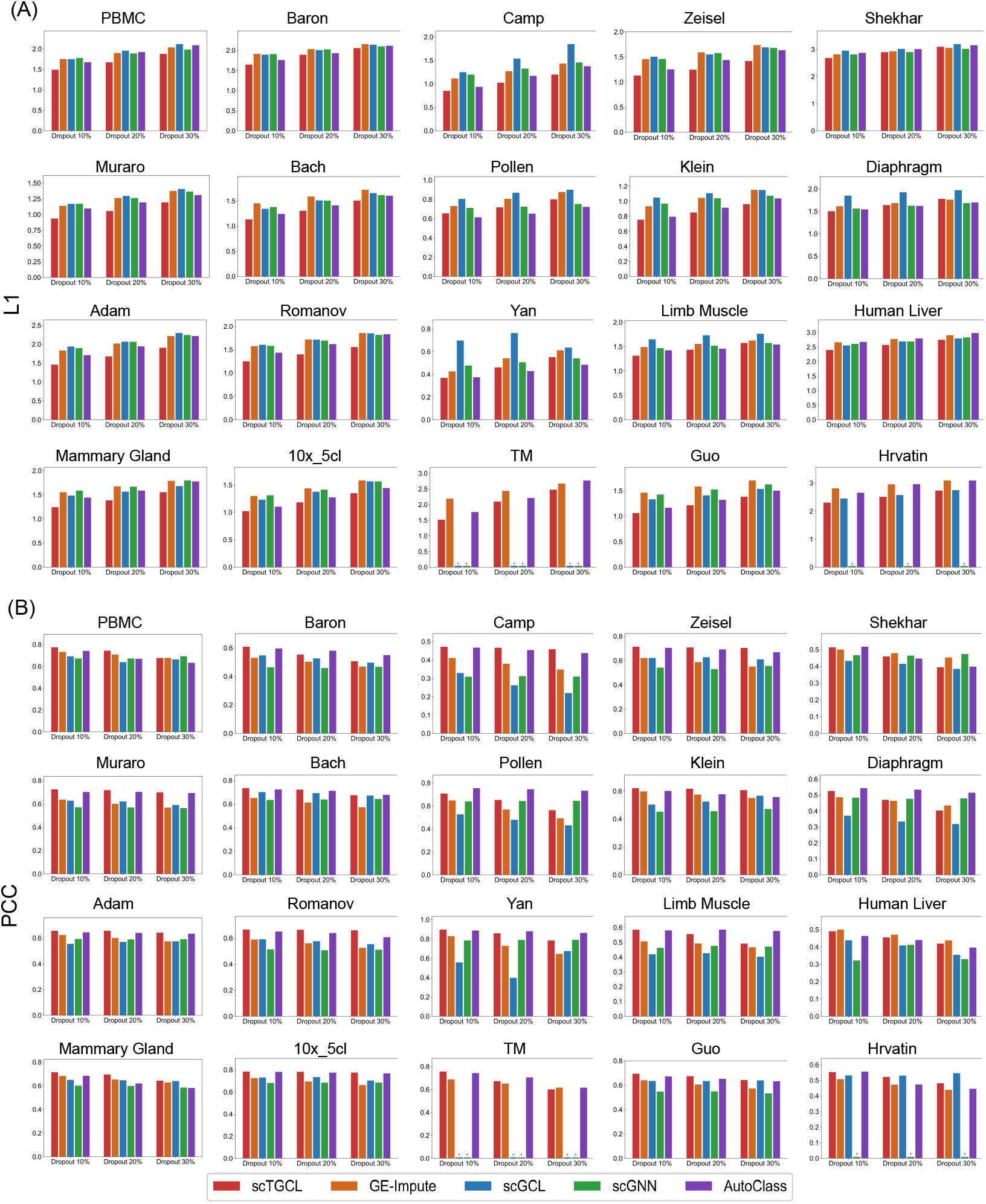
Imputation performance comparison on twenty real scRNA-seq datasets at 10%, 20%, and 30% artificial dropout rates. Both metrics are computed only on the entries that were artificially masked. (A) L1 distance (lower is better). (B) PCC (higher is better).

Figure 4 presents the L1 distance and PCC comparisons across all three dropout levels. As shown in Figure 4(A), scTGCL consistently achieves the lowest L1 distance among all compared methods at each dropout rate, indicating that the imputed expression values remain closest to the ground-truth data. Notably, the performance advantage of scTGCL widens as the dropout rate increases from 10% to 30%, demonstrating its robustness under high-sparsity conditions. Figure 4(B) further confirms this trend: scTGCL attains the highest PCC values across all dropout settings, reflecting strong correlation between recovered and original gene expression profiles. Among the baseline models, AutoClass achieves the second best performance overall, demonstrating that its deep autoencoder-based architecture generalizes well to expression recovery. Interestingly, despite incorporating graph contrastive learning, scGCL does not achieve competitive imputation performance, suggesting that its contrastive objective is not sufficiently aligned with dropout recovery and that graph contrastive learning alone does not guarantee strong imputation. This further highlights that joint optimization of contrastive, reconstruction, and imputation losses is what enables effective utilization of the scTGCL framework for expression recovery. GE-Impute and scGNN display moderate imputation accuracy with greater variability across datasets. It is worth noting that both scGCL and scGNN are computationally heavy models that could not be executed on large-scale datasets such as TM and Hrvatin due to memory constraints, and their results are therefore absent for those datasets. We also compared the whole reconstructed matrix with the original matrix by computing the PCC over all entries at 20%, 40%, and 60% dropout (Supplementary Figure S3). Here, scTGCL achieves the highest PCC on every dataset at 40% and 60% dropout, while AutoClass is slightly higher at 20% dropout. The advantage of scTGCL grows as dropout increases, which again shows that it is most useful when dropout is severe.

These results demonstrate that the joint optimization of imputation and contrastive learning objectives in scTGCL not only recovers dropout events accurately but also preserves the global transcriptomic structure essential for downstream biological analysis.

### 3.2. Ablation Study

To analyze the contribution of each core component of scTGCL, we conduct a comprehensive ablation study on all twenty real scRNA-seq datasets. Each model variant is evaluated ten times per dataset and the averaged results are reported in Figure 5.

**Fig 5.**
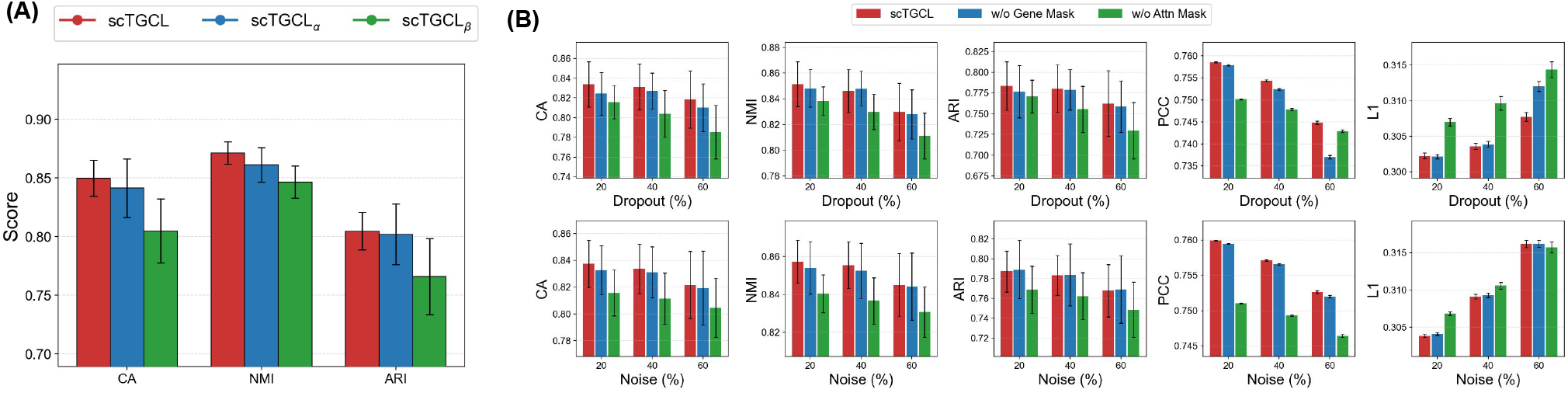
Performance comparison of scTGCL on twenty datasets. (A) CA, NMI, and ARI for full scTGCL, scTGCL_*α*_ (KNN graph based encoder), and scTGCL_*β*_ (no contrastive loss). (B) CA, NMI, ARI, PCC, and L1 for full scTGCL, scTGCL without gene masking (w/o Gene Mask) and scTGCL without attention masking (w/o Attn Mask) under 20%, 40%, and 60% dropout (top) and noise (bottom).

The central architectural innovation of scTGCL is its Transformer-based encoder, which employs multi-head self-attention to construct adaptive cell–cell graphs that capture diverse transcriptomic relationships without relying on predefined similarity measures. To validate its effectiveness, we define two ablated variants. scTGCL_*α*_ replaces the multi-head attention-based graph construction with a standard *k*-nearest-neighbour (KNN) graph, while keeping all other components unchanged. scTGCL_*β*_ retains the Transformer-based encoder but removes the graph contrastive learning loss, training the model with reconstruction and imputation objectives only. Figure 5(A) reports the average CA, NMI, and ARI across all twenty datasets. scTGCL consistently outperforms scTGCL_*α*_ on all three metrics, confirming that the adaptive attention-based graph construction captures biologically richer cell–cell relationships than a fixed KNN structure. The performance drop of scTGCL_*β*_ is substantially larger, demonstrating that graph contrastive learning is critical for learning discriminative and noise-resilient representations. Per-dataset results for this experiment are provided in Supplementary Table S1.

A second key component of scTGCL is its dual augmentation strategy, comprising random gene masking at the feature level and random attention masking on the self-attention matrices. Both mechanisms are designed to improve model robustness against dropout events and technical noise. To quantify their individual contributions, we evaluate two additional variants: w/o Gene Mask, which disables gene masking while retaining attention masking, and w/o Attn Mask, which disables attention masking while retaining gene masking. All three variants are tested under artificially imposed dropout rates of 20%, 40%, and 60%, and independently under noise levels of 20%, 40%, and 60%, with performance measured by CA, NMI, ARI, PCC, and L1. As shown in Figure 5(B), removing gene masking generally causes a modest performance decline, most clearly under artificial dropout, indicating that it contributes useful regularization by simulating expression dropout during training. In contrast, removing attention masking leads to substantially larger degradation in both clustering accuracy and imputation quality, particularly at higher dropout and noise rates, demonstrating that attention masking is the more critical of the two mechanisms for handling transcriptomic noise and sparsity. Per-dataset results for this experiment are provided in Supplementary Table S2 - Table S4.

Taken together, these results confirm that each component of scTGCL contributes positively to the overall performance, and their joint optimization is essential for achieving accurate, robust, and biologically meaningful clustering and imputation results.

### 3.3. Effect of Highly Variable Gene Selection

The selection of highly variable genes (HVGs) is a critical preprocessing step in scRNA-seq analysis, as it directly influences the quality of learned representations and downstream clustering performance. To investigate the robustness of scTGCL with respect to gene selection, we systematically evaluate its performance by varying the number of HVGs selected using the Scanpy framework. Specifically, we consider five different HVG settings: 500, 1500, 2500, 3500, and 4500 genes, and report CA, NMI and ARI across twenty real scRNA-seq datasets. Each model was evaluated ten times per HVG setting and the average is reported.

As shown in Figure 6, the clustering performance of scTGCL exhibits a slightly dependency on the number of selected HVGs. Using a small number of genes (e.g., 500 HVGs) leads to lower median scores and higher variance across datasets, indicating insufficient transcriptomic information. Performance improves consistently as the number of HVGs increases, reaching its peak at 2500 HVGs, where CA, ARI, and NMI achieve the highest median values with relatively low variance. This suggests that 2500 HVGs provide an optimal balance between capturing informative biological signals and suppressing noise. When the number of HVGs is further increased to 3500 or 4500, performance slightly degrades or becomes more variable, likely due to the inclusion of less informative or noisy genes. Overall, this analysis demonstrates that scTGCL is robust to HVG selection and that selecting approximately 2500 HVGs yields the most stable and optimal clustering performance across diverse datasets.

**Fig 6.**
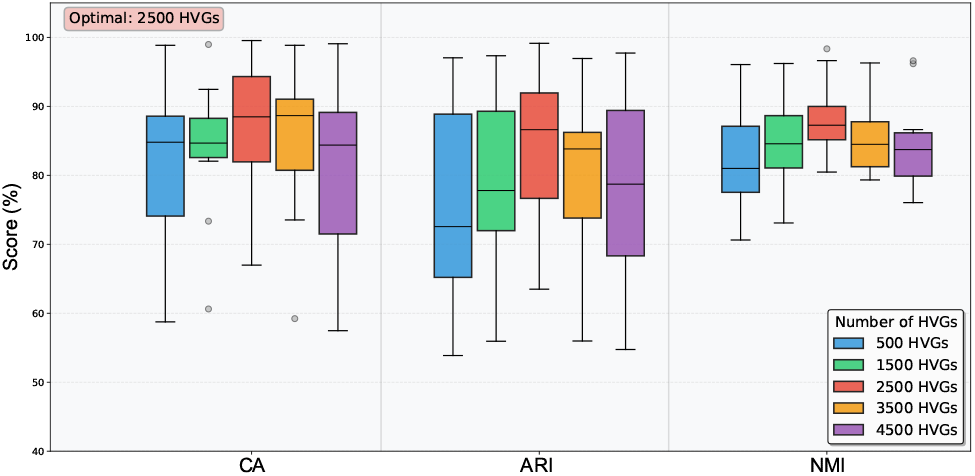
Effect of the number of selected highly variable genes (HVGs) on clustering performance. Boxplots summarize CA, ARI, and NMI scores across twenty scRNA-seq datasets for different HVG selections.

### 3.4. Robustness Analysis

To evaluate the robustness of the proposed scTGCL, we generated multiple simulated scRNA-seq datasets using the Splatter package [63] under different noise and complexity settings. We compared our model with nine baseline methods using CA, NMI and ARI metrics. All results are averaged over ten independent runs. To analyze the effect of dropout, we first varied the dropout midpoint parameter over the values 0.5, 1.0, 1.5, and 2.5, using datasets with 5000 cells, 2500 genes and 8 cell types. In Splatter, the probability that a gene is dropped out in a cell follows a logistic function of the gene’s mean expression, and this parameter sets the midpoint of this function. With the negative dropout shape used in our simulations, a larger midpoint makes dropout more frequent, so the simulated data become sparser as this value increases. As shown in Figure 7(A), scTGCL maintains the highest CA, NMI and ARI even when dropout becomes severe. Although scSimGCL also performs well, scTGCL consistently achieves slightly higher scores. This indicates that the proposed model can effectively learn meaningful representations even with large amounts of missing gene expression values. We further evaluated performance using different differential expression scale values (0.2, 0.25, 0.3, 0.35).

**Fig 7.**
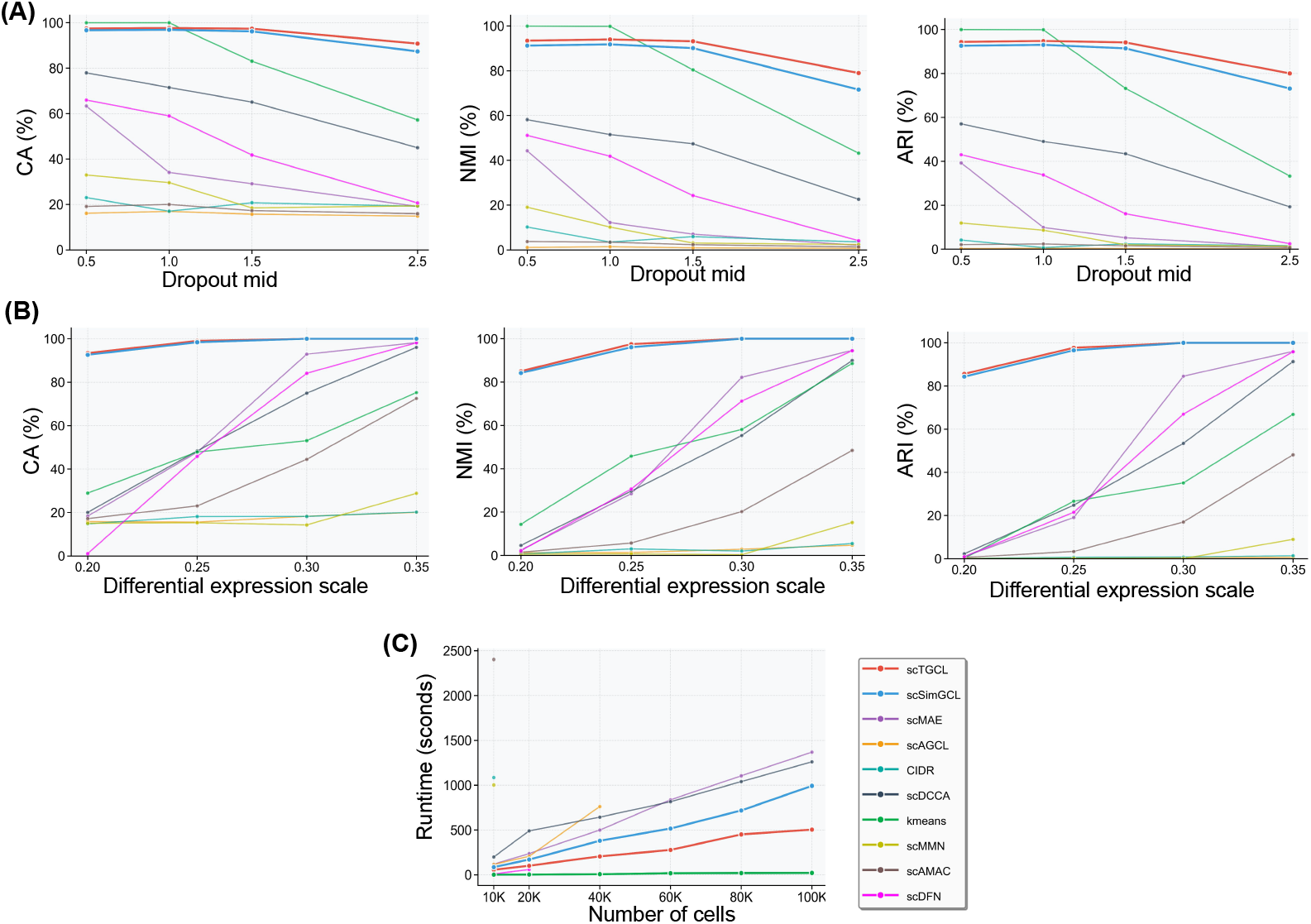
Robustness evaluation on simulated datasets. (A) Performance under different values of the Splatter dropout midpoint parameter (dropout.mid); larger values produce more dropout. (B) Performance under different differential expression scale. (C) Runtime comparison on datasets of increasing size.

Lower differential expression scale produces less separable cell populations, making clustering difficult. From Figure 7(B), scTGCL clearly outperforms all baselines in low differential expression scale settings, showing that it can distinguish subtle biological differences between cell groups. Finally, we measured runtime on datasets ranging from 10K to 100K cells. As shown in Figure 7(C), some methods (scAMAC, scMMN) fail due to memory limits, and others only run on small datasets. K-means is fastest but less accurate, while scTGCL achieves the second lowest runtime and successfully handles large datasets. This demonstrates that scTGCL is both efficient and scalable for large-scale scRNA-seq analysis.

### 3.5. Biological Analysis

To verify that the clusters found by scTGCL reflect real biological cell populations, we perform a biological analysis on the Muraro dataset [44], comprising cell type annotation, KEGG enrichment analysis, and marker gene identification.

For cell type annotation, scTGCL’s latent embeddings are partitioned into nine clusters with *k*-means, and COSG [64] is used to find the top 100 differentially expressed genes of each cluster. We then compare these genes with a curated marker panel for the nine known Muraro cell types, taken from the CellMarker database [65] and the literature, compute the overlap ratio with each type, and assign the cluster to the type with the highest ratio. As reported in Supplementary Figure S4, eight of the nine predicted clusters are confidently annotated as ductal, gamma, beta, mesenchymal, acinar, delta, alpha, and endothelial cells, with overlap ratios between 0.33 and 1.0, and each assignment agrees with the dominant ground truth label of its cluster. Cluster 8, however, shows no overlap with any panel; as it has only sixteen cells and its leading genes CD53, CD300A, and PTPRC resemble immune rather than pancreatic markers, we suspect it is a rare or mixed subpopulation and leave it unannotated as Unknown 8.

To explore the biological meaning of these genes, we perform KEGG [66] enrichment analysis through Enrichr [67], with the top enriched pathways of each cluster shown in Figure 8(A). The enriched pathways clearly differ across clusters, and the most significant pathway of each cluster matches the established physiology of the corresponding cell type, each of which has been independently confirmed in prior experimental studies. The beta cell cluster is most strongly enriched in ‘Maturity onset diabetes of the young’ (driven by INS, PDX1, MAFA, SLC2A2 and MNX1) together with ‘Insulin secretion’, mirroring the central role of beta cell transcription factors and the insulin secretory machinery in this monogenic diabetes, as established by Fajans and Bell [68]. The acinar cluster is dominated by ‘Pancreatic secretion’, its single most significant pathway across all clusters, reflecting the exocrine synthesis and secretion of digestive zymogens that defines acinar cells [69]. The ductal cluster is enriched in ‘Bile secretion’ through CA2, SLC4A4, CFTR and AQP1, consistent with the CFTR and bicarbonate based fluid secretion machinery shared by pancreatic duct cells and the biliary epithelium [70]. The alpha cell cluster is enriched in ‘Regulation of lipolysis in adipocytes’, in line with the well documented lipolytic action of glucagon, the hormone secreted by alpha cells [71]. The delta cell cluster is enriched in ‘Neuroactive ligand receptor interaction’ (including SST, GHSR, GHRH and LEPR), consistent with the broad G protein coupled receptor repertoire through which delta cells regulate somatostatin secretion [72]. The mesenchymal cluster is enriched in the collagen rich ‘Protein digestion and absorption’ and ‘ECM receptor interaction’ pathways, matching the role of pancreatic stellate and mesenchymal cells as the principal producers of extracellular matrix and collagen proteins [73]. Finally, the endothelial cluster is enriched in ‘Rap1 signaling’ and ‘Fluid shear stress and atherosclerosis’, pathways characteristic of vascular endothelium and its mechanotransduction of blood flow [74]. The consistency of these strongest, cell type specific pathways with independent experimental evidence further supports the plausibility of the annotations above.

**Fig 8.**
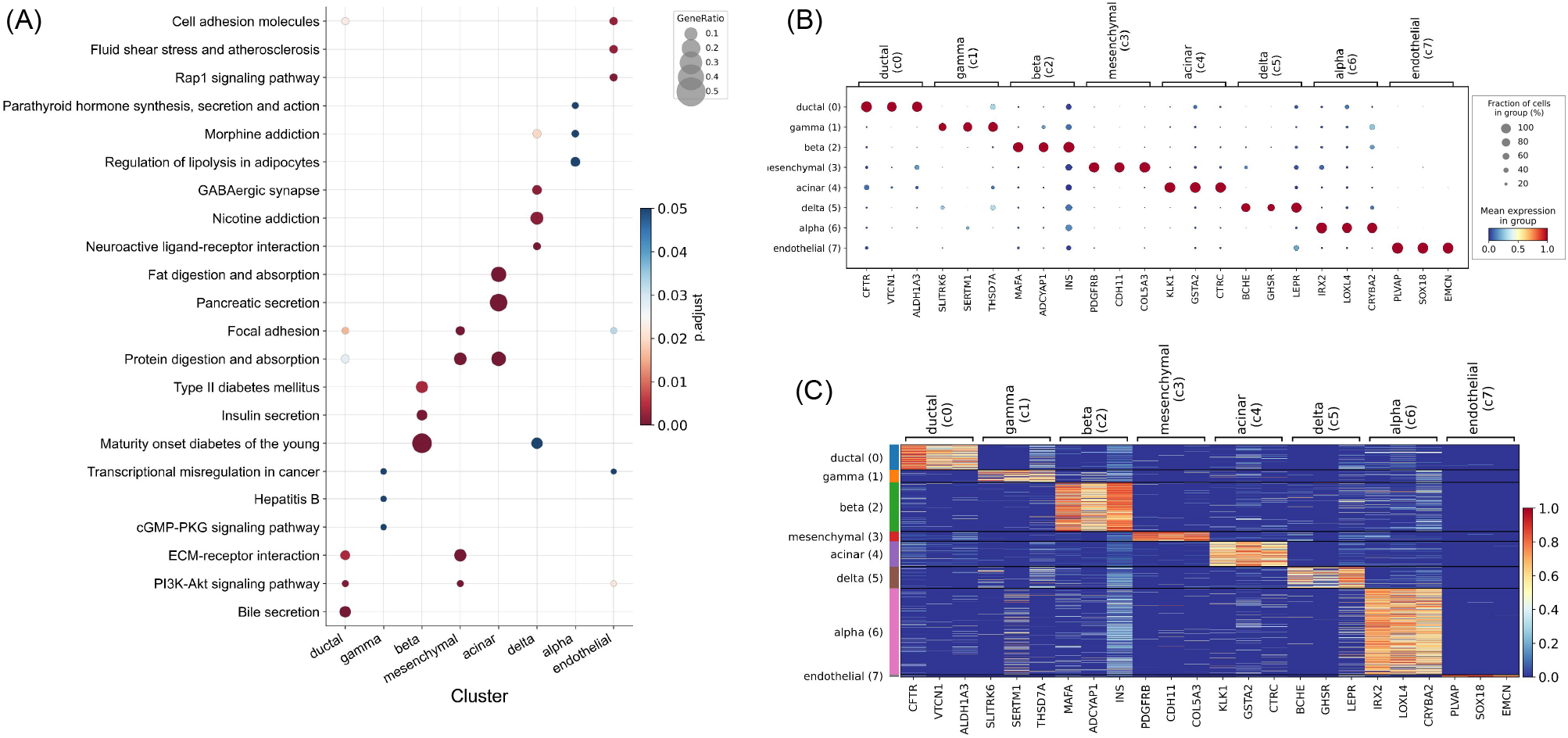
Biological analysis of scTGCL on the Muraro human pancreas dataset. (A) Top-10 KEGG-enriched pathways (ranked by *−* log_10_ adjusted *p*-value) for the COSG marker genes of each of the eight predicted clusters; cluster identities inferred from marker-gene overlap with known Muraro cell types are given in horizontal axis. (B) Dot plot of the top-3 COSG marker genes per cluster across the eight annotated clusters; dot size encodes the fraction of expressing cells and color encodes mean scaled expression. (C) Single-cell heat map of the same 24 marker genes (standard-scaled per gene), with cells grouped by predicted cluster and color bar on the left shows cell range.

In addition, we use COSG to identify the top three marker genes of each cluster and display their expression as a dot plot and a heat map, shown in Figure 8(B) and (C). Both show a clear diagonal pattern in which the marker genes of each cluster are expressed at high levels mainly within that cluster and remain low elsewhere, indicating that scTGCL’s clusters are transcriptionally specific and distinct. Comparing these genes with the CellMarker database [65] shows that most match the recorded markers of the corresponding cell type; for example, the alpha cell markers we identify, IRX2, LOXL4, and CRYBA2, are also documented there. Altogether, these findings show that scTGCL’s clustering results are biologically meaningful and well suited to downstream analyses such as cell annotation and marker gene discovery, valuable for understanding disease mechanisms.

### 3.6. Computational Efficiency

Modern scRNA-seq studies often profile tens of thousands of cells, so a clustering method must train quickly and fit within limited GPU memory. We therefore recorded the training time and the peak GPU memory of every method on all twenty datasets, ordered from the smallest (Yan, 90 cells) to the largest (TM, 54,865 cells) (Figure 9).

**Fig 9.**
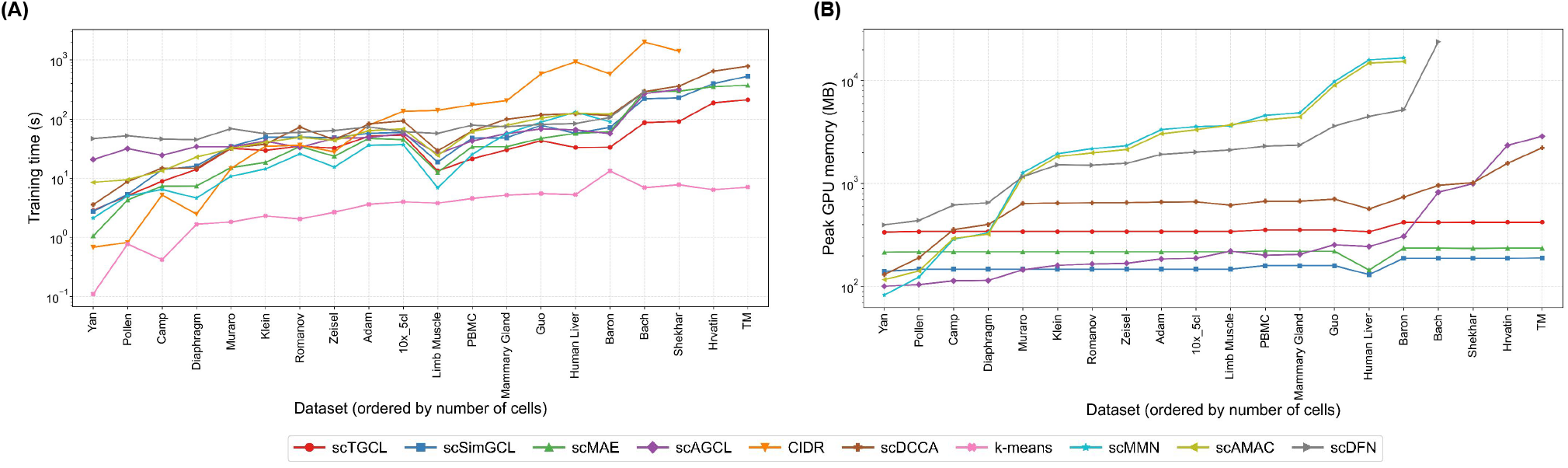
Computational efficiency of scTGCL and competing methods on twenty real scRNA-seq datasets, ordered by number of cells. (A) Training time in seconds (log scale). (B) Peak GPU memory during training in MB (log scale). Missing points indicate that a method could not complete training on that dataset. k-means and CIDR run on the CPU and are therefore not shown in (B).

Figure 9(A) shows the training time. On small datasets, some lightweight methods such as CIDR and scMAE finish faster than scTGCL. As the number of cells grows, however, this changes. On every dataset with more than about 4,000 cells, scTGCL needs the least training time among all deep learning methods, and on the largest datasets, Bach, Shekhar, Hrvatin, and TM, it finishes in roughly half the time of the next fastest one. k-means is the fastest method on every dataset, but its clustering accuracy is much lower (Figure 2), so its speed comes at a large cost in quality.

Figure 9(B) shows the peak GPU memory. The memory use of scTGCL stays almost constant, at about 330 to 420 MB, from the smallest to the largest dataset. This is because scTGCL computes attention only within each mini-batch, so its memory depends on the batch size rather than on the total number of cells. scSimGCL and scMAE use less memory than scTGCL, so scTGCL lies in the middle of the compared methods. In contrast, the memory of scAGCL and scDCCA keeps rising with dataset size, and that of scMMN, scAMAC, and scDFN grows rapidly to more than 10 GB. scMMN, scAMAC, scDFN, and CIDR could not complete training on the largest datasets. Overall, scTGCL combines the shortest training time among deep learning methods on large datasets with a moderate and stable memory requirement, which makes it practical for large-scale scRNA-seq analysis.

## 4. Conclusion

In this paper, we proposed scTGCL, a novel Transformer-based graph contrastive learning framework for clustering single-cell RNA-seq data. The model introduces a multi-head attention mechanism to learn adaptive cell–cell graphs directly from gene expression data without relying on predefined similarity measures. By integrating feature-level augmentation through gene masking and graph-level augmentation through edge dropping, scTGCL generates robust augmented views that effectively simulate dropout events and structural uncertainty inherent in scRNA-seq data. A symmetric contrastive loss is employed to maximize agreement between original and augmented representations while preserving discriminative information, jointly optimized with reconstruction and imputation objectives to maintain biological interpretability.

Extensive experiments on twenty real scRNA-seq datasets demonstrated that scTGCL achieves the highest average performance among nine state-of-the-art baseline methods in clustering accuracy, normalized mutual information, and adjusted Rand index. The ablation study confirmed the essential contribution of each architectural component, while robustness analysis on simulated datasets showed that scTGCL maintains high performance under varying dropout rates and differential expression levels. Furthermore, the model exhibits superior computational efficiency, achieving substantially lower runtime on large-scale datasets compared with existing graph-based and contrastive learning approaches. Hyperparameter analysis further validated the stability of scTGCL across different batch sizes and loss weight configurations.

Overall, scTGCL provides a simple, efficient, and scalable solution for robust single-cell clustering. By combining Transformer-based graph learning with contrastive representation learning, the proposed framework offers a promising direction for future developments in single-cell transcriptomic analysis and related biomedical applications. Nevertheless, scTGCL has two main limitations. First, the model learns only from gene expression values and does not use known relationships between genes. In future work, we plan to integrate gene–gene interaction networks from databases such as STRING or Reactome, for example as prior knowledge that guides how the model combines information from related genes, which could further improve representation quality and clustering accuracy. Second, scTGCL does not explicitly correct for batch effects, which arise when cells are profiled in different experiments, laboratories, or sequencing platforms and remain a major challenge in single-cell clustering. Because cells from different batches can look different for technical rather than biological reasons, we plan to make scTGCL robust to batch effects, for example by adding batch information to the contrastive objective so that the model groups cells by cell type rather than by batch.

## 5. Author contributions

Md. Shoaib Abdullah Khan performed formal analysis, data curation, simulations and software development and wrote the original draft of the manuscript as well as contributed to review and editing. Md. Mahir Faisal contributed to formal analysis, data curation, simulations, software development and manuscript review and editing. Md. Humayun Kabir conceptualized the study, supervised the project, contributed to formal analysis and participated in writing the original draft as well as review and editing of the manuscript.

## Notes

### Competing Interest Statement

The authors have declared no competing interest.

### Summary of Updates

In this revised version, the manuscript has been substantially updated in response to the reviewers' comments. The abstract has been revised to state the research gap and the main findings more accurately. The introduction has been restructured to enrich the literature review with recent graph contrastive and Transformer-based methods, and the novelty of scTGCL has been redefined and highlighted as attention masking for contrastive view generation. In the methods, the mathematical notation and equations have been corrected and the batch-wise computation of attention has been clarified. In the results, the clustering comparison has been redrawn with standard deviations across runs, the performance discussion has been rewritten with more precise and accurate claims, the imputation evaluation has been clarified, the ablation study has been refined, and the robustness analysis now describes the simulation parameters correctly. A new computational efficiency analysis of training time and GPU memory replaces the earlier runtime table. The conclusion now discusses the limitations of the model together with concrete future directions, and the supplementary material has been reorganized with an additional figure and properly titled tables and figures.

