## Supplementary material for "scTGCL: A Transformer-Based Graph Contrastive Learning Approach for Efficiently Clustering Single-Cell RNA-seq Data": Supplimentary material

**Contents**

**1. Supplementary Tables**

Table S1. Clustering performance of scTGCL and its ablated variants on twenty scRNA-seq datasets

Table S2. Robustness of scTGCL under artificial dropout and noise

Table S3. Robustness of scTGCL without gene masking under artificial dropout and noise

Table S4. Robustness of scTGCL without attention masking under artificial dropout and noise

**2. Supplementary Figures**

Figure S1. Clustering performance measured by ARI on twenty real scRNA-seq datasets

Figure S2. Effect of batch size and hyperparameters on clustering performance

Figure S3. Imputation performance on the whole reconstructed expression matrix

Figure S4. Cell type annotation of scTGCL clusters on the Muraro dataset

**1. Supplementary Tables**

**Table S1. Clustering performance of scTGCL and its ablated variants on twenty scRNA-seq datasets.** CA, NMI, and ARI of the full scTGCL model, of scTGCL-α, which replaces the learned attention graph with a fixed k-nearest-neighbour graph, and of scTGCL-β, which removes the contrastive loss. These per-dataset results correspond to Figure 5A of the main text.

| **Dataset** | **scTGCL** | | | **scTGCL-β** | | | **scTGCL-α** | | |
| --- | --- | --- | --- | --- | --- | --- | --- | --- | --- |
|  | **CA** | **NMI** | **ARI** | **CA** | **NMI** | **ARI** | **CA** | **NMI** | **ARI** |
| adam | 0.8790 | 0.8350 | 0.7695 | 0.9180 | 0.8538 | 0.8260 | 0.8590 | 0.8337 | 0.7640 |
| pollen | 0.9541 | 0.9780 | 0.9729 | 0.9213 | 0.9588 | 0.9269 | 0.9377 | 0.9655 | 0.9485 |
| pbmc | 0.8402 | 0.7958 | 0.7758 | 0.8376 | 0.7956 | 0.7832 | 0.8304 | 0.7923 | 0.7675 |
| baron | 0.6318 | 0.8116 | 0.5638 | 0.5518 | 0.7508 | 0.5243 | 0.5880 | 0.7806 | 0.5332 |
| muraro | 0.9435 | 0.9079 | 0.9241 | 0.9431 | 0.8934 | 0.9230 | 0.9172 | 0.8699 | 0.8996 |
| bach | 0.7935 | 0.8197 | 0.8044 | 0.8129 | 0.8308 | 0.8355 | 0.8699 | 0.8471 | 0.8957 |
| klein | 0.8452 | 0.8469 | 0.8030 | 0.8445 | 0.8091 | 0.8001 | 0.9099 | 0.8743 | 0.8746 |
| camp | 0.8013 | 0.8987 | 0.7588 | 0.6321 | 0.7889 | 0.6045 | 0.8026 | 0.8641 | 0.7312 |
| zeisel | 0.9052 | 0.7920 | 0.8273 | 0.8473 | 0.7977 | 0.7698 | 0.8988 | 0.7858 | 0.8254 |
| shekhar | 0.7252 | 0.8534 | 0.5589 | 0.6015 | 0.7937 | 0.5388 | 0.7397 | 0.8666 | 0.6719 |
| diaphragm | 0.9931 | 0.9833 | 0.9903 | 0.9966 | 0.9871 | 0.9912 | 0.9931 | 0.9737 | 0.9815 |
| romanov | 0.8929 | 0.7460 | 0.7678 | 0.7338 | 0.6358 | 0.5985 | 0.8873 | 0.7390 | 0.7637 |
| yan | 0.7322 | 0.8608 | 0.6878 | 0.7000 | 0.8564 | 0.6212 | 0.7444 | 0.8656 | 0.6806 |
| limb muscle | 0.9853 | 0.9699 | 0.9693 | 0.7844 | 0.8652 | 0.7249 | 0.9138 | 0.9054 | 0.9164 |
| human_liver | 0.7556 | 0.8245 | 0.7063 | 0.7519 | 0.8308 | 0.7184 | 0.7287 | 0.8043 | 0.6896 |
| mammary gland | 0.9619 | 0.9474 | 0.9309 | 0.9405 | 0.9224 | 0.9368 | 0.9146 | 0.9108 | 0.9194 |
| 10x_5cl | 1.0000 | 1.0000 | 1.0000 | 1.0000 | 1.0000 | 1.0000 | 1.0000 | 1.0000 | 1.0000 |
| TM | 0.6314 | 0.8213 | 0.6653 | 0.6027 | 0.8173 | 0.6046 | 0.5751 | 0.8056 | 0.5698 |
| guo | 0.8690 | 0.8665 | 0.8088 | 0.7866 | 0.8344 | 0.7131 | 0.8587 | 0.8640 | 0.8048 |
| hrvatin | 0.9333 | 0.9394 | 0.9286 | 0.8904 | 0.9066 | 0.8777 | 0.8580 | 0.8760 | 0.8021 |

**Table S2. Robustness of scTGCL under artificial dropout and noise.** Clustering (CA, NMI, ARI) and imputation (PCC, L1) performance of the full scTGCL model on twenty real scRNA-seq datasets when 20%, 40%, or 60% of the expression entries are corrupted by artificial dropout or by noise. These per-dataset results correspond to Figure 5B of the main text.

| **Dataset** | **Corruption (%)** | **Dropout** | | | | | **Noise** | | | | |
| --- | --- | --- | --- | --- | --- | --- | --- | --- | --- | --- | --- |
|  |  | **CA** | **NMI** | **ARI** | **PCC** | **L1** | **CA** | **NMI** | **ARI** | **PCC** | **L1** |
| adam | 20% | 0.8891 | 0.8523 | 0.7937 | 0.8046 | 0.2660 | 0.9167 | 0.8729 | 0.8218 | 0.8050 | 0.2667 |
|  | 40% | 0.8954 | 0.8366 | 0.7793 | 0.8029 | 0.2676 | 0.9104 | 0.8595 | 0.8106 | 0.8033 | 0.2713 |
|  | 60% | 0.8087 | 0.7619 | 0.6686 | 0.7994 | 0.2707 | 0.9030 | 0.8425 | 0.7937 | 0.8005 | 0.2778 |
| pollen | 20% | 0.9443 | 0.9737 | 0.9669 | 0.7996 | 0.2719 | 0.9475 | 0.9769 | 0.9705 | 0.8004 | 0.2712 |
|  | 40% | 0.9443 | 0.9741 | 0.9681 | 0.7980 | 0.2737 | 0.9475 | 0.9769 | 0.9705 | 0.7992 | 0.2735 |
|  | 60% | 0.9344 | 0.9590 | 0.9526 | 0.7943 | 0.2787 | 0.9443 | 0.9737 | 0.9669 | 0.7971 | 0.2773 |
| pbmc | 20% | 0.8281 | 0.7837 | 0.7563 | 0.7638 | 0.2670 | 0.8290 | 0.7838 | 0.7588 | 0.7638 | 0.2701 |
|  | 40% | 0.8224 | 0.7740 | 0.7475 | 0.7617 | 0.2654 | 0.8212 | 0.7748 | 0.7425 | 0.7609 | 0.2747 |
|  | 60% | 0.8113 | 0.7509 | 0.7286 | 0.7559 | 0.2643 | 0.8035 | 0.7526 | 0.7055 | 0.7565 | 0.2807 |
| baron | 20% | 0.5898 | 0.7717 | 0.5437 | 0.7237 | 0.2737 | 0.6084 | 0.7835 | 0.5702 | 0.7257 | 0.2775 |
|  | 40% | 0.5998 | 0.7708 | 0.5490 | 0.7153 | 0.2745 | 0.5856 | 0.7737 | 0.5476 | 0.7203 | 0.2851 |
|  | 60% | 0.6112 | 0.7583 | 0.5515 | 0.6921 | 0.2816 | 0.6008 | 0.7801 | 0.5549 | 0.7118 | 0.2950 |
| muraro | 20% | 0.9304 | 0.8849 | 0.8822 | 0.7909 | 0.3284 | 0.9642 | 0.9147 | 0.9315 | 0.7914 | 0.3285 |
|  | 40% | 0.9600 | 0.9032 | 0.9237 | 0.7891 | 0.3303 | 0.9525 | 0.9056 | 0.9264 | 0.7897 | 0.3323 |
|  | 60% | 0.9092 | 0.8752 | 0.8635 | 0.7853 | 0.3342 | 0.9416 | 0.9002 | 0.9233 | 0.7869 | 0.3377 |
| bach | 20% | 0.7836 | 0.8082 | 0.8040 | 0.7874 | 0.3000 | 0.7849 | 0.8101 | 0.8055 | 0.7881 | 0.3032 |
|  | 40% | 0.7818 | 0.8054 | 0.8030 | 0.7839 | 0.2999 | 0.7829 | 0.8098 | 0.8050 | 0.7857 | 0.3082 |
|  | 60% | 0.7810 | 0.8012 | 0.8008 | 0.7734 | 0.3074 | 0.7822 | 0.8084 | 0.8048 | 0.7819 | 0.3150 |
| klein | 20% | 0.9507 | 0.9004 | 0.9208 | 0.6921 | 0.5263 | 0.9467 | 0.9037 | 0.9175 | 0.6929 | 0.5256 |
|  | 40% | 0.9437 | 0.8906 | 0.9118 | 0.6904 | 0.5279 | 0.9408 | 0.8933 | 0.9086 | 0.6924 | 0.5266 |
|  | 60% | 0.9456 | 0.8755 | 0.9044 | 0.6870 | 0.5307 | 0.9298 | 0.8755 | 0.8904 | 0.6914 | 0.5282 |
| camp | 20% | 0.7718 | 0.8050 | 0.6444 | 0.8116 | 0.4041 | 0.7885 | 0.8212 | 0.6697 | 0.8122 | 0.4023 |
|  | 40% | 0.7679 | 0.8003 | 0.6397 | 0.8103 | 0.4080 | 0.7795 | 0.8124 | 0.6577 | 0.8114 | 0.4049 |
|  | 60% | 0.7885 | 0.8217 | 0.6760 | 0.8074 | 0.4152 | 0.7692 | 0.8022 | 0.6440 | 0.8100 | 0.4090 |
| zeisel | 20% | 0.8639 | 0.7465 | 0.7726 | 0.7956 | 0.3094 | 0.8815 | 0.7558 | 0.7950 | 0.7964 | 0.3099 |
|  | 40% | 0.8802 | 0.7483 | 0.7907 | 0.7932 | 0.3112 | 0.8842 | 0.7612 | 0.8018 | 0.7946 | 0.3141 |
|  | 60% | 0.8715 | 0.7353 | 0.7709 | 0.7878 | 0.3159 | 0.8825 | 0.7572 | 0.7985 | 0.7916 | 0.3203 |
| shekhar | 20% | 0.6702 | 0.8005 | 0.4904 | 0.5675 | 0.3258 | 0.6536 | 0.7991 | 0.4811 | 0.5710 | 0.3353 |
|  | 40% | 0.6340 | 0.7650 | 0.4738 | 0.5585 | 0.3154 | 0.6457 | 0.8007 | 0.4995 | 0.5683 | 0.3382 |
|  | 60% | 0.6029 | 0.7209 | 0.4586 | 0.5350 | 0.3045 | 0.6478 | 0.7895 | 0.4761 | 0.5640 | 0.3423 |
| diaphragm | 20% | 0.9851 | 0.9437 | 0.9620 | 0.6487 | 0.3001 | 0.9828 | 0.9352 | 0.9560 | 0.6507 | 0.2986 |
|  | 40% | 0.9839 | 0.9407 | 0.9591 | 0.6439 | 0.3036 | 0.9701 | 0.9261 | 0.9528 | 0.6478 | 0.3020 |
|  | 60% | 0.9805 | 0.9309 | 0.9503 | 0.6347 | 0.3107 | 0.9586 | 0.9212 | 0.9526 | 0.6431 | 0.3072 |
| romanov | 20% | 0.8243 | 0.7079 | 0.6910 | 0.7705 | 0.3259 | 0.7983 | 0.6817 | 0.6454 | 0.7712 | 0.3272 |
|  | 40% | 0.8537 | 0.7280 | 0.7299 | 0.7680 | 0.3266 | 0.8572 | 0.7206 | 0.7198 | 0.7694 | 0.3310 |
|  | 60% | 0.8544 | 0.7213 | 0.7236 | 0.7625 | 0.3293 | 0.7712 | 0.6636 | 0.6135 | 0.7666 | 0.3365 |
| yan | 20% | 0.6667 | 0.8488 | 0.5807 | 0.9272 | 0.2132 | 0.6667 | 0.8488 | 0.5807 | 0.9275 | 0.2128 |
|  | 40% | 0.6667 | 0.8488 | 0.5807 | 0.9264 | 0.2152 | 0.6667 | 0.8488 | 0.5807 | 0.9270 | 0.2142 |
|  | 60% | 0.6556 | 0.8371 | 0.5622 | 0.9246 | 0.2188 | 0.6667 | 0.8488 | 0.5807 | 0.9263 | 0.2163 |
| limb muscle | 20% | 0.9862 | 0.9516 | 0.9604 | 0.6837 | 0.2870 | 0.9890 | 0.9604 | 0.9679 | 0.6854 | 0.2860 |
|  | 40% | 0.9679 | 0.9437 | 0.9526 | 0.6793 | 0.2910 | 0.9908 | 0.9667 | 0.9727 | 0.6822 | 0.2905 |
|  | 60% | 0.9523 | 0.9339 | 0.9437 | 0.6706 | 0.2968 | 0.9523 | 0.9354 | 0.9451 | 0.6770 | 0.2969 |
| human_liver | 20% | 0.7049 | 0.7886 | 0.6581 | 0.6791 | 0.3096 | 0.7303 | 0.8138 | 0.6855 | 0.6850 | 0.3141 |
|  | 40% | 0.7076 | 0.7789 | 0.6585 | 0.6641 | 0.3115 | 0.7423 | 0.8208 | 0.6897 | 0.6771 | 0.3258 |
|  | 60% | 0.6651 | 0.7317 | 0.5974 | 0.6325 | 0.3163 | 0.7201 | 0.8071 | 0.6732 | 0.6648 | 0.3410 |
| mammary gland | 20% | 0.9151 | 0.8940 | 0.9156 | 0.7589 | 0.2311 | 0.9242 | 0.9010 | 0.9269 | 0.7591 | 0.2338 |
|  | 40% | 0.9133 | 0.8877 | 0.9117 | 0.7558 | 0.2319 | 0.9153 | 0.8957 | 0.9163 | 0.7551 | 0.2407 |
|  | 60% | 0.9193 | 0.8835 | 0.9121 | 0.7487 | 0.2353 | 0.9322 | 0.9022 | 0.9319 | 0.7487 | 0.2500 |
| 10x_5cl | 20% | 1.0000 | 1.0000 | 1.0000 | 0.7781 | 0.4851 | 1.0000 | 1.0000 | 1.0000 | 0.7786 | 0.4845 |
|  | 40% | 1.0000 | 1.0000 | 1.0000 | 0.7768 | 0.4874 | 1.0000 | 1.0000 | 1.0000 | 0.7778 | 0.4866 |
|  | 60% | 1.0000 | 1.0000 | 1.0000 | 0.7741 | 0.4912 | 1.0000 | 1.0000 | 1.0000 | 0.7765 | 0.4898 |
| TM | 20% | 0.6005 | 0.8076 | 0.6217 | 0.8781 | 0.1415 | 0.5669 | 0.8037 | 0.5433 | 0.8784 | 0.1470 |
|  | 40% | 0.5887 | 0.8022 | 0.5999 | 0.8736 | 0.1480 | 0.5487 | 0.7973 | 0.5046 | 0.8720 | 0.1660 |
|  | 60% | 0.5579 | 0.7818 | 0.5575 | 0.8637 | 0.1603 | 0.5179 | 0.7909 | 0.4740 | 0.8621 | 0.1874 |
| guo | 20% | 0.8733 | 0.8605 | 0.8297 | 0.7758 | 0.2811 | 0.8488 | 0.8566 | 0.8040 | 0.7765 | 0.2827 |
|  | 40% | 0.8379 | 0.8485 | 0.7800 | 0.7734 | 0.2828 | 0.8119 | 0.8409 | 0.7467 | 0.7747 | 0.2881 |
|  | 60% | 0.8393 | 0.8433 | 0.7783 | 0.7679 | 0.2880 | 0.8160 | 0.8456 | 0.7582 | 0.7718 | 0.2954 |
| hrvatin | 20% | 0.8966 | 0.8970 | 0.8725 | 0.7340 | 0.1978 | 0.9199 | 0.9251 | 0.9089 | 0.7389 | 0.2002 |
|  | 40% | 0.8759 | 0.8739 | 0.8427 | 0.7224 | 0.1997 | 0.9194 | 0.9240 | 0.9081 | 0.7329 | 0.2086 |
|  | 60% | 0.8784 | 0.8712 | 0.8437 | 0.6994 | 0.2042 | 0.8932 | 0.9025 | 0.8677 | 0.7234 | 0.2193 |

**Table S3. Robustness of scTGCL without gene masking under artificial dropout and noise.** Clustering (CA, NMI, ARI) and imputation (PCC, L1) performance of scTGCL without gene masking (w/o Gene Mask) on twenty real scRNA-seq datasets when 20%, 40%, or 60% of the expression entries are corrupted by artificial dropout or by noise. These per-dataset results correspond to Figure 5B of the main text.

| **Dataset** | **Corruption (%)** | **Dropout** | | | | | **Noise** | | | | |
| --- | --- | --- | --- | --- | --- | --- | --- | --- | --- | --- | --- |
|  |  | **CA** | **NMI** | **ARI** | **PCC** | **L1** | **CA** | **NMI** | **ARI** | **PCC** | **L1** |
| adam | 20% | 0.8850 | 0.8461 | 0.8032 | 0.8027 | 0.2679 | 0.9022 | 0.8610 | 0.8262 | 0.8032 | 0.2682 |
|  | 40% | 0.8757 | 0.8246 | 0.7811 | 0.8011 | 0.2696 | 0.9142 | 0.8644 | 0.8369 | 0.8017 | 0.2719 |
|  | 60% | 0.8003 | 0.7344 | 0.6591 | 0.7979 | 0.2724 | 0.9107 | 0.8547 | 0.8272 | 0.7993 | 0.2772 |
| pollen | 20% | 0.9180 | 0.9552 | 0.9152 | 0.7990 | 0.2711 | 0.9246 | 0.9596 | 0.9312 | 0.7999 | 0.2700 |
|  | 40% | 0.9246 | 0.9596 | 0.9312 | 0.7972 | 0.2738 | 0.9311 | 0.9641 | 0.9472 | 0.7989 | 0.2722 |
|  | 60% | 0.9213 | 0.9511 | 0.9300 | 0.7941 | 0.2780 | 0.9213 | 0.9569 | 0.9288 | 0.7969 | 0.2758 |
| pbmc | 20% | 0.8295 | 0.7811 | 0.7534 | 0.7628 | 0.2684 | 0.8346 | 0.7900 | 0.7657 | 0.7630 | 0.2695 |
|  | 40% | 0.8221 | 0.7695 | 0.7465 | 0.7611 | 0.2690 | 0.8304 | 0.7832 | 0.7610 | 0.7607 | 0.2733 |
|  | 60% | 0.8173 | 0.7553 | 0.7329 | 0.7574 | 0.2703 | 0.8113 | 0.7588 | 0.7180 | 0.7571 | 0.2784 |
| baron | 20% | 0.5870 | 0.7687 | 0.5455 | 0.7266 | 0.2747 | 0.5839 | 0.7679 | 0.5469 | 0.7288 | 0.2775 |
|  | 40% | 0.5883 | 0.7714 | 0.5463 | 0.7183 | 0.2765 | 0.5963 | 0.7744 | 0.5518 | 0.7226 | 0.2852 |
|  | 60% | 0.5935 | 0.7517 | 0.5328 | 0.6970 | 0.2835 | 0.6004 | 0.7706 | 0.5532 | 0.7130 | 0.2952 |
| muraro | 20% | 0.9539 | 0.9090 | 0.9268 | 0.7919 | 0.3267 | 0.9666 | 0.9175 | 0.9358 | 0.7925 | 0.3265 |
|  | 40% | 0.9553 | 0.9109 | 0.9277 | 0.7899 | 0.3291 | 0.9482 | 0.9078 | 0.9249 | 0.7909 | 0.3298 |
|  | 60% | 0.9153 | 0.8895 | 0.9105 | 0.7858 | 0.3336 | 0.9496 | 0.8994 | 0.9188 | 0.7883 | 0.3346 |
| bach | 20% | 0.7811 | 0.8052 | 0.8034 | 0.7894 | 0.2989 | 0.7822 | 0.8067 | 0.8042 | 0.7903 | 0.3022 |
|  | 40% | 0.7804 | 0.8043 | 0.8037 | 0.7854 | 0.2999 | 0.7798 | 0.8050 | 0.8025 | 0.7875 | 0.3075 |
|  | 60% | 0.7780 | 0.7996 | 0.8007 | 0.7730 | 0.3102 | 0.7787 | 0.8020 | 0.8009 | 0.7829 | 0.3150 |
| klein | 20% | 0.8283 | 0.8328 | 0.7885 | 0.6908 | 0.5276 | 0.8276 | 0.8291 | 0.7875 | 0.6916 | 0.5265 |
|  | 40% | 0.8265 | 0.8235 | 0.7852 | 0.6890 | 0.5300 | 0.8287 | 0.8305 | 0.7875 | 0.6911 | 0.5278 |
|  | 60% | 0.8022 | 0.7857 | 0.7580 | 0.6855 | 0.5349 | 0.8140 | 0.8116 | 0.7759 | 0.6902 | 0.5296 |
| camp | 20% | 0.7641 | 0.7984 | 0.6427 | 0.8112 | 0.4041 | 0.7910 | 0.8233 | 0.6786 | 0.8118 | 0.4021 |
|  | 40% | 0.7769 | 0.8197 | 0.6719 | 0.8099 | 0.4079 | 0.7603 | 0.8003 | 0.6455 | 0.8110 | 0.4050 |
|  | 60% | 0.7333 | 0.7816 | 0.6261 | 0.8073 | 0.4150 | 0.7603 | 0.7986 | 0.6473 | 0.8098 | 0.4090 |
| zeisel | 20% | 0.8522 | 0.7441 | 0.7605 | 0.7933 | 0.3115 | 0.8835 | 0.7652 | 0.8066 | 0.7940 | 0.3118 |
|  | 40% | 0.8859 | 0.7595 | 0.7999 | 0.7911 | 0.3133 | 0.8775 | 0.7601 | 0.7954 | 0.7923 | 0.3156 |
|  | 60% | 0.8386 | 0.7270 | 0.7384 | 0.7865 | 0.3173 | 0.8742 | 0.7524 | 0.7893 | 0.7895 | 0.3212 |
| shekhar | 20% | 0.6780 | 0.8239 | 0.5281 | 0.5608 | 0.3174 | 0.7281 | 0.8476 | 0.6038 | 0.5653 | 0.3368 |
|  | 40% | 0.6628 | 0.7944 | 0.5104 | 0.5398 | 0.3018 | 0.7142 | 0.8295 | 0.5554 | 0.5630 | 0.3391 |
|  | 60% | 0.6128 | 0.7339 | 0.4701 | 0.4484 | 0.3260 | 0.6611 | 0.8164 | 0.5283 | 0.5593 | 0.3423 |
| diaphragm | 20% | 0.9253 | 0.8841 | 0.9169 | 0.6483 | 0.3006 | 0.9253 | 0.8753 | 0.8947 | 0.6500 | 0.2994 |
|  | 40% | 0.9483 | 0.9101 | 0.9472 | 0.6436 | 0.3045 | 0.9253 | 0.8665 | 0.8725 | 0.6469 | 0.3032 |
|  | 60% | 0.9598 | 0.9184 | 0.9496 | 0.6353 | 0.3111 | 0.9207 | 0.8707 | 0.9065 | 0.6420 | 0.3089 |
| romanov | 20% | 0.8551 | 0.7203 | 0.7296 | 0.7686 | 0.3279 | 0.8589 | 0.7345 | 0.7449 | 0.7694 | 0.3282 |
|  | 40% | 0.8915 | 0.7586 | 0.7828 | 0.7663 | 0.3290 | 0.8329 | 0.7158 | 0.7111 | 0.7678 | 0.3317 |
|  | 60% | 0.8614 | 0.7431 | 0.7500 | 0.7612 | 0.3321 | 0.8582 | 0.7349 | 0.7411 | 0.7651 | 0.3369 |
| yan | 20% | 0.6667 | 0.8488 | 0.5807 | 0.9237 | 0.2183 | 0.6667 | 0.8488 | 0.5807 | 0.9238 | 0.2174 |
|  | 40% | 0.6667 | 0.8488 | 0.5807 | 0.9231 | 0.2198 | 0.6667 | 0.8488 | 0.5807 | 0.9232 | 0.2190 |
|  | 60% | 0.6667 | 0.8488 | 0.5807 | 0.9216 | 0.2230 | 0.6667 | 0.8488 | 0.5807 | 0.9223 | 0.2212 |
| limb muscle | 20% | 0.9945 | 0.9818 | 0.9861 | 0.6827 | 0.2867 | 0.9936 | 0.9780 | 0.9826 | 0.6844 | 0.2857 |
|  | 40% | 0.9954 | 0.9846 | 0.9883 | 0.6784 | 0.2900 | 0.9954 | 0.9840 | 0.9871 | 0.6814 | 0.2899 |
|  | 60% | 0.9936 | 0.9789 | 0.9831 | 0.6698 | 0.2972 | 0.9743 | 0.9668 | 0.9719 | 0.6766 | 0.2960 |
| human_liver | 20% | 0.7483 | 0.8113 | 0.6974 | 0.6834 | 0.3016 | 0.7349 | 0.8070 | 0.6839 | 0.6908 | 0.3106 |
|  | 40% | 0.7301 | 0.7825 | 0.6693 | 0.6611 | 0.3035 | 0.7563 | 0.8186 | 0.7070 | 0.6811 | 0.3241 |
|  | 60% | 0.7243 | 0.7593 | 0.6722 | 0.6050 | 0.3187 | 0.7150 | 0.7972 | 0.6565 | 0.6660 | 0.3416 |
| mammary gland | 20% | 0.8433 | 0.8570 | 0.7936 | 0.7581 | 0.2334 | 0.8725 | 0.8747 | 0.8507 | 0.7584 | 0.2353 |
|  | 40% | 0.8939 | 0.8890 | 0.8673 | 0.7552 | 0.2350 | 0.8890 | 0.8874 | 0.8636 | 0.7546 | 0.2417 |
|  | 60% | 0.9171 | 0.8905 | 0.9066 | 0.7488 | 0.2385 | 0.9072 | 0.8903 | 0.8996 | 0.7485 | 0.2504 |
| 10x_5cl | 20% | 1.0000 | 1.0000 | 1.0000 | 0.7775 | 0.4856 | 1.0000 | 1.0000 | 1.0000 | 0.7781 | 0.4845 |
|  | 40% | 1.0000 | 1.0000 | 1.0000 | 0.7762 | 0.4882 | 1.0000 | 1.0000 | 1.0000 | 0.7774 | 0.4865 |
|  | 60% | 1.0000 | 1.0000 | 1.0000 | 0.7735 | 0.4929 | 1.0000 | 1.0000 | 1.0000 | 0.7763 | 0.4895 |
| TM | 20% | 0.5772 | 0.8033 | 0.6247 | 0.8783 | 0.1425 | 0.5935 | 0.8098 | 0.6412 | 0.8788 | 0.1475 |
|  | 40% | 0.5676 | 0.7936 | 0.5710 | 0.8730 | 0.1509 | 0.5574 | 0.8007 | 0.5790 | 0.8722 | 0.1665 |
|  | 60% | 0.5413 | 0.7745 | 0.5265 | 0.8601 | 0.1694 | 0.5170 | 0.7891 | 0.4680 | 0.8619 | 0.1878 |
| guo | 20% | 0.8715 | 0.8586 | 0.8281 | 0.7741 | 0.2836 | 0.8733 | 0.8605 | 0.8309 | 0.7749 | 0.2832 |
|  | 40% | 0.8743 | 0.8587 | 0.8388 | 0.7718 | 0.2869 | 0.8732 | 0.8610 | 0.8322 | 0.7734 | 0.2877 |
|  | 60% | 0.8703 | 0.8531 | 0.8322 | 0.7670 | 0.2931 | 0.8436 | 0.8499 | 0.7963 | 0.7709 | 0.2941 |
| hrvatin | 20% | 0.9256 | 0.9319 | 0.9064 | 0.7340 | 0.1940 | 0.9115 | 0.9261 | 0.8865 | 0.7394 | 0.1996 |
|  | 40% | 0.8717 | 0.8962 | 0.8277 | 0.7169 | 0.1984 | 0.9435 | 0.9455 | 0.9333 | 0.7334 | 0.2080 |
|  | 60% | 0.8570 | 0.8837 | 0.8113 | 0.6653 | 0.2231 | 0.9016 | 0.9143 | 0.8710 | 0.7236 | 0.2188 |

**Table S4. Robustness of scTGCL without attention masking under artificial dropout and noise.** Clustering (CA, NMI, ARI) and imputation (PCC, L1) performance of scTGCL without attention masking (w/o Attn Mask) on twenty real scRNA-seq datasets when 20%, 40%, or 60% of the expression entries are corrupted by artificial dropout or by noise. These per-dataset results correspond to Figure 5B of the main text.

| **Dataset** | **Corruption (%)** | **Dropout** | | | | | **Noise** | | | | |
| --- | --- | --- | --- | --- | --- | --- | --- | --- | --- | --- | --- |
|  |  | **CA** | **NMI** | **ARI** | **PCC** | **L1** | **CA** | **NMI** | **ARI** | **PCC** | **L1** |
| adam | 20% | 0.8858 | 0.8526 | 0.7998 | 0.8024 | 0.2669 | 0.9183 | 0.8729 | 0.8255 | 0.8030 | 0.2671 |
|  | 40% | 0.8445 | 0.8022 | 0.7258 | 0.8009 | 0.2688 | 0.8492 | 0.8206 | 0.7416 | 0.8019 | 0.2701 |
|  | 60% | 0.7669 | 0.7277 | 0.6369 | 0.7979 | 0.2730 | 0.8656 | 0.8166 | 0.7396 | 0.8000 | 0.2745 |
| pollen | 20% | 0.9475 | 0.9650 | 0.9575 | 0.7998 | 0.2704 | 0.9377 | 0.9618 | 0.9432 | 0.8005 | 0.2697 |
|  | 40% | 0.9311 | 0.9592 | 0.9412 | 0.7981 | 0.2733 | 0.9311 | 0.9580 | 0.9366 | 0.7993 | 0.2723 |
|  | 60% | 0.9115 | 0.9509 | 0.9032 | 0.7948 | 0.2782 | 0.9213 | 0.9510 | 0.9174 | 0.7973 | 0.2761 |
| pbmc | 20% | 0.8366 | 0.7819 | 0.7641 | 0.7608 | 0.2705 | 0.8408 | 0.7869 | 0.7767 | 0.7610 | 0.2712 |
|  | 40% | 0.8339 | 0.7732 | 0.7623 | 0.7595 | 0.2715 | 0.8346 | 0.7774 | 0.7627 | 0.7595 | 0.2736 |
|  | 60% | 0.8235 | 0.7612 | 0.7482 | 0.7568 | 0.2728 | 0.8221 | 0.7606 | 0.7348 | 0.7571 | 0.2770 |
| baron | 20% | 0.5328 | 0.7269 | 0.4869 | 0.6925 | 0.2884 | 0.5349 | 0.7363 | 0.4897 | 0.6934 | 0.2886 |
|  | 40% | 0.5354 | 0.7177 | 0.4642 | 0.6894 | 0.2911 | 0.5471 | 0.7344 | 0.4960 | 0.6916 | 0.2910 |
|  | 60% | 0.5251 | 0.6993 | 0.4496 | 0.6835 | 0.2943 | 0.5397 | 0.7263 | 0.4868 | 0.6885 | 0.2950 |
| muraro | 20% | 0.9445 | 0.9028 | 0.9251 | 0.7907 | 0.3276 | 0.9609 | 0.9130 | 0.9305 | 0.7914 | 0.3268 |
|  | 40% | 0.9544 | 0.9126 | 0.9323 | 0.7887 | 0.3300 | 0.9379 | 0.9040 | 0.9268 | 0.7900 | 0.3298 |
|  | 60% | 0.9266 | 0.8905 | 0.9173 | 0.7845 | 0.3351 | 0.9266 | 0.8936 | 0.8891 | 0.7878 | 0.3343 |
| bach | 20% | 0.7837 | 0.8091 | 0.8088 | 0.7780 | 0.3108 | 0.7829 | 0.8090 | 0.8087 | 0.7787 | 0.3111 |
|  | 40% | 0.7807 | 0.8047 | 0.8053 | 0.7763 | 0.3127 | 0.7820 | 0.8078 | 0.8078 | 0.7776 | 0.3130 |
|  | 60% | 0.7780 | 0.7972 | 0.7999 | 0.7726 | 0.3161 | 0.7792 | 0.8064 | 0.8059 | 0.7758 | 0.3158 |
| klein | 20% | 0.8202 | 0.7451 | 0.7378 | 0.6860 | 0.5303 | 0.8224 | 0.7444 | 0.7384 | 0.6864 | 0.5293 |
|  | 40% | 0.8254 | 0.7488 | 0.7436 | 0.6850 | 0.5312 | 0.8221 | 0.7446 | 0.7377 | 0.6860 | 0.5299 |
|  | 60% | 0.8213 | 0.7439 | 0.7388 | 0.6826 | 0.5348 | 0.8217 | 0.7404 | 0.7359 | 0.6854 | 0.5306 |
| camp | 20% | 0.7231 | 0.7589 | 0.5830 | 0.8076 | 0.4107 | 0.7205 | 0.7529 | 0.5756 | 0.8084 | 0.4091 |
|  | 40% | 0.7000 | 0.7552 | 0.5768 | 0.8063 | 0.4140 | 0.7128 | 0.7428 | 0.5602 | 0.8076 | 0.4108 |
|  | 60% | 0.6987 | 0.7435 | 0.5668 | 0.8037 | 0.4207 | 0.7141 | 0.7441 | 0.5620 | 0.8065 | 0.4137 |
| zeisel | 20% | 0.8892 | 0.7699 | 0.8068 | 0.7900 | 0.3129 | 0.8845 | 0.7764 | 0.8032 | 0.7907 | 0.3126 |
|  | 40% | 0.8469 | 0.7482 | 0.7550 | 0.7879 | 0.3154 | 0.8958 | 0.7780 | 0.8133 | 0.7895 | 0.3158 |
|  | 60% | 0.8519 | 0.7447 | 0.7583 | 0.7838 | 0.3199 | 0.8915 | 0.7747 | 0.8111 | 0.7873 | 0.3206 |
| shekhar | 20% | 0.5349 | 0.7193 | 0.4482 | 0.5477 | 0.3416 | 0.5455 | 0.7258 | 0.4427 | 0.5488 | 0.3423 |
|  | 40% | 0.5129 | 0.6989 | 0.4204 | 0.5451 | 0.3428 | 0.5406 | 0.7239 | 0.4404 | 0.5477 | 0.3440 |
|  | 60% | 0.4752 | 0.6551 | 0.3949 | 0.5402 | 0.3427 | 0.5251 | 0.7098 | 0.4314 | 0.5459 | 0.3467 |
| diaphragm | 20% | 0.9828 | 0.9352 | 0.9560 | 0.6418 | 0.2982 | 0.9828 | 0.9352 | 0.9560 | 0.6430 | 0.2973 |
|  | 40% | 0.9828 | 0.9352 | 0.9560 | 0.6388 | 0.3016 | 0.9828 | 0.9352 | 0.9560 | 0.6414 | 0.2997 |
|  | 60% | 0.9828 | 0.9365 | 0.9561 | 0.6327 | 0.3076 | 0.9828 | 0.9352 | 0.9560 | 0.6388 | 0.3034 |
| romanov | 20% | 0.8548 | 0.7268 | 0.7318 | 0.7623 | 0.3339 | 0.7955 | 0.6926 | 0.6775 | 0.7632 | 0.3334 |
|  | 40% | 0.7972 | 0.7098 | 0.6889 | 0.7604 | 0.3358 | 0.8575 | 0.7278 | 0.7360 | 0.7621 | 0.3357 |
|  | 60% | 0.7733 | 0.6875 | 0.6588 | 0.7562 | 0.3394 | 0.8520 | 0.7182 | 0.7258 | 0.7602 | 0.3392 |
| yan | 20% | 0.6667 | 0.8488 | 0.5807 | 0.9237 | 0.2215 | 0.6667 | 0.8488 | 0.5807 | 0.9242 | 0.2199 |
|  | 40% | 0.6667 | 0.8488 | 0.5807 | 0.9229 | 0.2235 | 0.6667 | 0.8488 | 0.5807 | 0.9236 | 0.2212 |
|  | 60% | 0.6556 | 0.8169 | 0.5329 | 0.9213 | 0.2269 | 0.6667 | 0.8488 | 0.5807 | 0.9231 | 0.2225 |
| limb muscle | 20% | 0.9927 | 0.9729 | 0.9776 | 0.6774 | 0.2860 | 0.9936 | 0.9760 | 0.9800 | 0.6788 | 0.2853 |
|  | 40% | 0.9908 | 0.9679 | 0.9721 | 0.6742 | 0.2882 | 0.9927 | 0.9729 | 0.9776 | 0.6768 | 0.2883 |
|  | 60% | 0.9917 | 0.9712 | 0.9747 | 0.6673 | 0.2934 | 0.9936 | 0.9760 | 0.9800 | 0.6734 | 0.2929 |
| human_liver | 20% | 0.6173 | 0.7522 | 0.5459 | 0.6516 | 0.3137 | 0.6575 | 0.7815 | 0.5988 | 0.6542 | 0.3133 |
|  | 40% | 0.5962 | 0.7424 | 0.5221 | 0.6454 | 0.3182 | 0.6133 | 0.7601 | 0.5385 | 0.6504 | 0.3213 |
|  | 60% | 0.5427 | 0.6955 | 0.4560 | 0.6327 | 0.3255 | 0.6301 | 0.7665 | 0.5660 | 0.6442 | 0.3319 |
| mammary gland | 20% | 0.8988 | 0.9015 | 0.9095 | 0.7550 | 0.2352 | 0.8734 | 0.8901 | 0.8602 | 0.7555 | 0.2350 |
|  | 40% | 0.8936 | 0.8850 | 0.8931 | 0.7525 | 0.2378 | 0.8986 | 0.9021 | 0.9092 | 0.7527 | 0.2400 |
|  | 60% | 0.8698 | 0.8719 | 0.8483 | 0.7477 | 0.2428 | 0.8943 | 0.8939 | 0.9035 | 0.7480 | 0.2473 |
| 10x_5cl | 20% | 1.0000 | 1.0000 | 1.0000 | 0.7761 | 0.4866 | 1.0000 | 1.0000 | 1.0000 | 0.7766 | 0.4855 |
|  | 40% | 1.0000 | 1.0000 | 1.0000 | 0.7749 | 0.4893 | 1.0000 | 1.0000 | 1.0000 | 0.7760 | 0.4871 |
|  | 60% | 0.9995 | 0.9981 | 0.9992 | 0.7725 | 0.4939 | 1.0000 | 1.0000 | 1.0000 | 0.7750 | 0.4898 |
| TM | 20% | 0.6067 | 0.8084 | 0.6325 | 0.8762 | 0.1437 | 0.5997 | 0.8110 | 0.6239 | 0.8766 | 0.1480 |
|  | 40% | 0.5937 | 0.8019 | 0.6130 | 0.8723 | 0.1501 | 0.5643 | 0.8029 | 0.5622 | 0.8707 | 0.1660 |
|  | 60% | 0.5435 | 0.7772 | 0.5346 | 0.8645 | 0.1605 | 0.5119 | 0.7878 | 0.4466 | 0.8614 | 0.1869 |
| guo | 20% | 0.8750 | 0.8594 | 0.8476 | 0.7715 | 0.2854 | 0.8753 | 0.8614 | 0.8466 | 0.7723 | 0.2847 |
|  | 40% | 0.8721 | 0.8596 | 0.8447 | 0.7695 | 0.2888 | 0.8769 | 0.8606 | 0.8479 | 0.7711 | 0.2885 |
|  | 60% | 0.8692 | 0.8511 | 0.8340 | 0.7655 | 0.2958 | 0.8772 | 0.8621 | 0.8504 | 0.7691 | 0.2940 |
| hrvatin | 20% | 0.9221 | 0.9288 | 0.9134 | 0.7124 | 0.2055 | 0.9234 | 0.9310 | 0.9150 | 0.7139 | 0.2063 |
|  | 40% | 0.9210 | 0.9260 | 0.9109 | 0.7075 | 0.2086 | 0.9227 | 0.9298 | 0.9142 | 0.7098 | 0.2132 |
|  | 60% | 0.9005 | 0.9042 | 0.8828 | 0.6979 | 0.2140 | 0.8778 | 0.9003 | 0.8501 | 0.7032 | 0.2220 |

**2. Supplementary Figures**

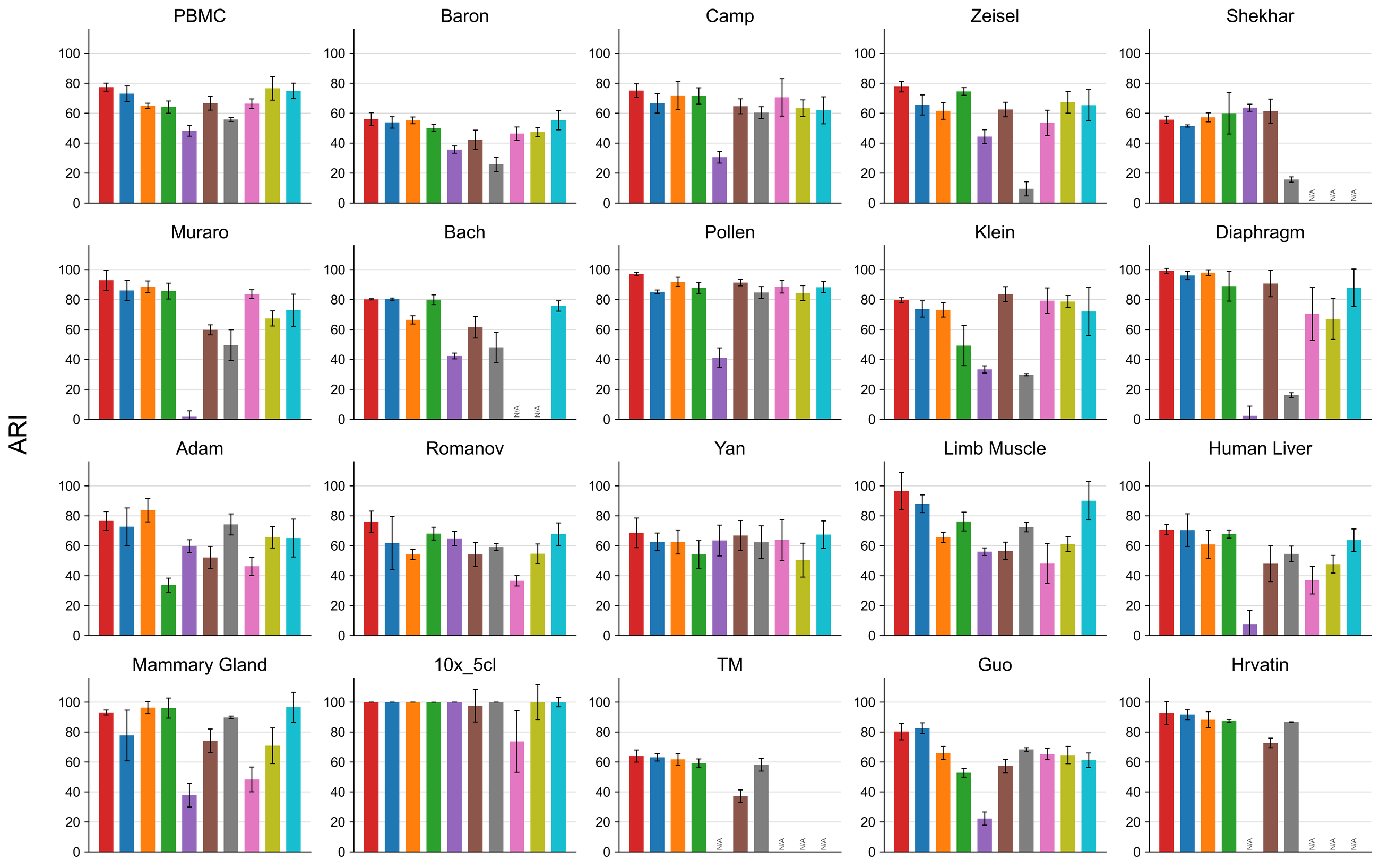

**Figure S1. Clustering performance measured by ARI on twenty real scRNA-seq datasets.** Each bar shows the mean adjusted Rand index (ARI, %) over ten independent runs with different random seeds, and the error bar shows the standard deviation across these runs. Methods are shown in the same order and colours as in Figure 2 of the main text: scTGCL (red), scSimGCL (blue), scMAE (orange), scAGCL (green), CIDR (purple), scDCCA (brown), k-means (grey), scMMN (pink), scAMAC (olive), and scDFN (cyan). N/A marks a method that could not be run on that dataset.

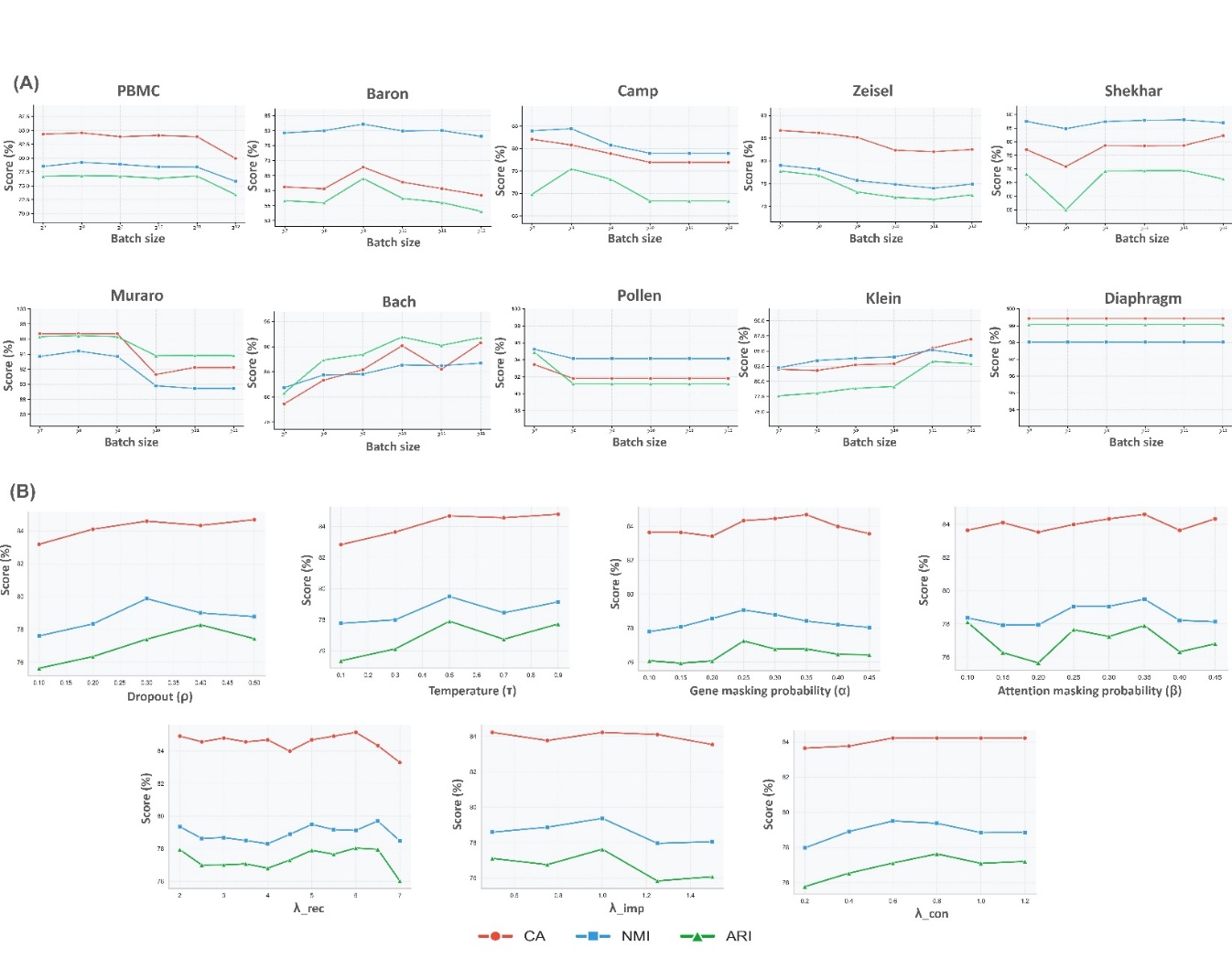

**Figure S2. Effect of batch size and hyperparameters on clustering performance.** (A) CA, NMI, and ARI of scTGCL with batch sizes from 128 (2^7) to 4096 (2^12) on ten real scRNA-seq datasets. (B) CA, NMI, and ARI on the PBMC dataset when varying the dropout rate (ρ), the contrastive temperature (τ), the gene masking probability (α), the attention masking probability (β), and the loss weights λ_rec, λ_imp, and λ_con, with all other settings fixed.

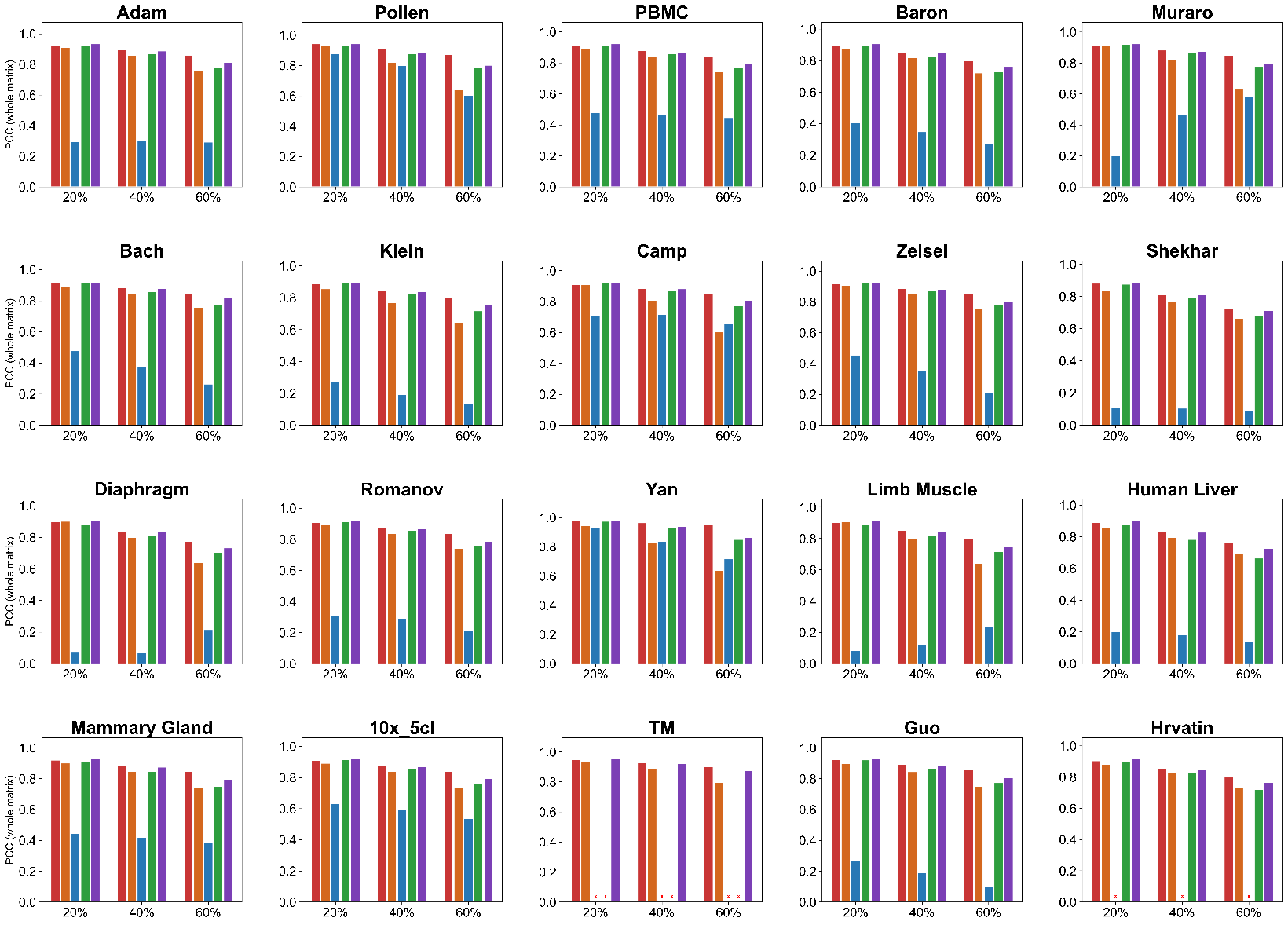

**Figure S3. Imputation performance on the whole reconstructed expression matrix.** Pearson correlation coefficient (PCC) between the whole reconstructed expression matrix and the original matrix for scTGCL (red), GE-Impute (orange), scGCL (blue), scGNN (green), and AutoClass (purple) on twenty real scRNA-seq datasets at 20%, 40%, and 60% artificial dropout. A red × marks a method that produced no result on that dataset.

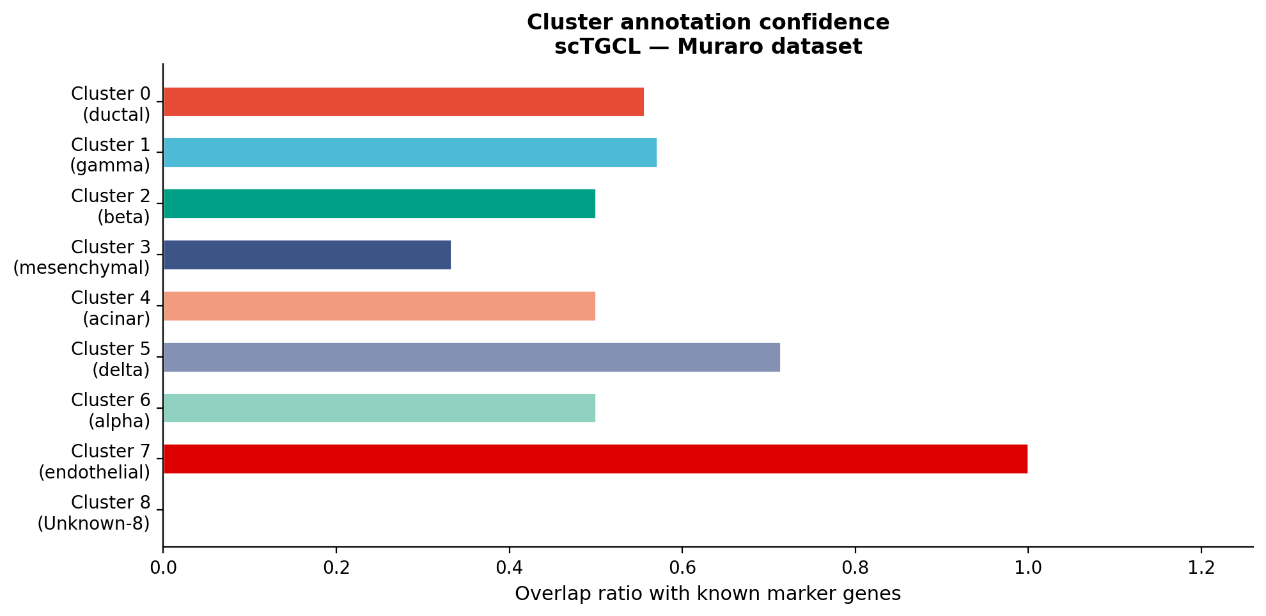

**Figure S4. Cell type annotation of scTGCL clusters on the Muraro dataset.** For each of the nine clusters found by scTGCL, the bar shows the overlap ratio between the top 100 COSG marker genes of the cluster and the curated marker panel of its assigned cell type. Cluster 8 shows no overlap with any known cell type and is labelled Unknown-8.
